# Differences in the co-distribution of the Cannabinoid Receptor-1, FAAH, and MAGL in the human and mouse brain could be limiting clinical translation of endocannabinoid outcomes in mouse models of Alzheimer disease

**DOI:** 10.64898/2025.12.10.693257

**Authors:** Ryan M. Heistad, Jennifer N.K. Nyarko, Paul R. Pennington, Huzaifa Saeed, Lais V.B. Gomes, Darrell D. Mousseau

## Abstract

Endocannabinoid system (ECS) outcomes in mouse models of Alzheimer disease (AD) do not always align with the clinical disease course. We compared the distribution of the Cannabinoid Receptor-1 (CB1R) and the enzymes, FAAH and MAGL, in autopsy AD brain samples as well as in the ‘J20’ (hAPP_Swe/Ind_) mouse model of AD. Both sources revealed several anti-CB1R immunoreactive species, *e.g*. one at 47-kDa (corresponding to the protein-coding sequence) and a reported putative splice variant at 37 kDa. We did not observe any changes in the mean expression in CB1R, FAAH or MAGL in the human samples, but did observe sex- and genotype-specific changes in the mouse brain. Regression analysis revealed strong sex- and *APOE* ε4-dependent associations among the CB1Rs as well as between CB1Rs and MAGL (but not FAAH) in human cortical (but not hippocampal) samples. In the J20 mouse, associations between CB1Rs were limited to hippocampal samples, whereas associations between CB1R and both MAGL and FAAH were observed in the cortex. A diagnosis of AD disrupted any associations in the human dataset, whereas in several instances, the associations were enhanced by the hAPP_Swe/Ind_ transgene. This inferred species-dependent regulation of the ECS could impact the clinical translational of ECS outcomes in preclinical models of AD pathology.

## Introduction

The endocannabinoid system (ECS) distributes throughout the brain [1] and, not surprisingly, it has been implicated in brain health and basic neurotransmission [2] as well as in processes that underpin mood and neuropsychiatric phenotypes [3] and degenerative diseases and therapies [4, 5]. The ECS involves two cannabinoid receptors (CB1R, CB2R). The CB1R is ubiquitously expressed in brain, whereas the CB2R, although being expressed at far lower levels, may have more (sub)regional or cell type-specific (and inducible) distribution [6]. This report will focus on the CB1R. The putative endogenous ECS ligands are N-arachidonoyl-ethanolamine (anandamide: AEA) [7] and 2-arachidonoyl glycerol (2-AG) [8] and a gas chromatography/mass spectrometry study of rapidly frozen rat brain was able to estimate 2-AG levels to be 174 times greater than those of AEA [9]. Fatty acid amide hydrolase (FAAH) and monoacylglycerol lipase (MAGL) are the primary enzymes responsible for AEA [10] and 2-AG [11] degradation, respectively. As estimating brain levels of AEA and 2-AG can be challenging [12], the expression of FAAH and MAGL are often used as proxies for the abundance of AEA and 2-AG.

ECS ligands are synthesized post-synaptically and act as retrograde neuromodulators [13]. The distribution of MAGL and the CB1R to pre-synaptic terminals is thought to allow MAGL to inactivate 2-AG within proximity of its site of action, whereas the distribution of FAAH to post-synaptic terminals is thought to allow regulation of AEA availability close to its site of synthesis [14–16]. This disparate distribution of enzymes and sites of ligand inactivation are now thought to provide distinct roles for the two ligands, with 2-AG acting as a retrograde neuromodulator and AEA underpinning tonic actions of the ECS and contributing to synaptic plasticity, potentially in a region-dependent manner [17]. The influence of ECS ligands *via* the CB1R on several neurotransmitter systems, particularly GABA and glutamate [18], and their modulation of transmitter release that indirectly affects long-term potentiation and depression [19], aligns well with the role of the ECS in memory and cognition [4]. Additional interactions with monoaminergic systems [20] implicates the ECS in mood and neuropsychiatric disorders [21–23]. Preclinical (animal) models often align with clinical evidence and support a role for the ECS in, for example, post-traumatic stress disorder [24], stress resilience [25], nausea and vomiting [26] as well as in anxiety, schizophrenia and depression [27–29]. While reports often extrapolate rodent-based CB1R-related data to their human orthologs, differences between animal models and humans must be considered. For example, a meta-analysis highlighted differences in CB1R affinity and regional distribution in the rat and human brain [30], while in the mouse brain, receptor binding [31] and, more recently, MALDI-mass spectrometry imaging [32], both find highest levels of 2-AG (which do not necessarily correlate with those of AEA) in the hypothalamus, whereas moderate levels are found in cortical and hindbrain structures, and the lowest levels are in the hippocampus. In the rat brain, levels of 2-AG and AEA are highest in the hippocampus, striatum, and brainstem and levels of 2-AG and AEA only correlate in regions that express the CBRs [33]. Of note, although relatively heterogeneously distributed throughout the neocortex of rat, monkey, and human species, CB1R immunoreactivity is least dense in layer 4 of cortical structures in the rat [34], but has the highest density in the same cortical layer of the human and monkey brain [35]. Differences in the expression patterns of components of the ECS extend to the periphery in mouse, rat, and monkey [36].

There is strong interest in defining the influence of the ECS during the course of diseases of the brain associated with aging, such as Alzheimer disease (AD) (see reviews [4, 37–39]). AD presents with behavioral, thinking, and cognitive deficits that reflect pathology centered on the hippocampus and associated structures [40], which appear to spread from pathology that is triggered across different cortical regions; such a phenomenon is associated with normal aging, but exacerbated in AD [41]. Interestingly, CB1R densities are relatively higher in these two regions [42]. Aside from advancing age [43], other risk factors, including biological sex [44] and genetics [45], are helping to identify modifiable events within the earlier stages of AD progression. By far, the majority of cases of AD present clinically after the age of 65 years and these late-onset cases of AD (LOAD) are of unknown cause, although markers of the disease can be identified as early as 25-30 years before symptoms appear [46, 47]. Cases of early-onset AD (EOAD) account for fewer than 10% of all cases of AD and are distinguished as non-mendelian (non-familial) cases (∼8%) or the rare cases that involve mutations in the *APP*, *PSEN1* or *PSEN2* genes (∼1-2%), all of which tend to increase levels of toxic β-amyloid (Aβ) length variants [46, 47]. Models support a protective role for the ECS that is lost or in deficit during the clinical course of AD. The response of the CB1R in AD is somewhat ambiguous, with both losses in density (*e.g*. [48, 49]) and no differences from age-matched controls (*e.g*. [50, 51]) being reported. Elsewhere, autoradiography of post-mortem tissues using selective CB1R radioligands [^3^H]CP55,940 [52] and [^125^I]SD-7015 [53] suggests CB1R density may actually undergo a bi-phasic response during the course of AD, with an increase in early Braak staging and then diminishing as Tau pathology is exacerbated, and may follow a subregion-specific response (*e.g*., within cortical layers or within hippocampal subfields). FAAH and MAGL may exert different effects during different stages of AD/dementia [54], which would alter endogenous ligand availability and CB1R number and functionality [55, 56]. One of the autoradiography studies referred to above [52] did not find any correlation between CB1R density and cognitive status, nor did it find any change in CB1R density in the nucleus basalis of Meynert, a region that is well known to be impacted during the earliest stages of AD [57]. In contrast, CB1R activity in the nucleus basalis of Meynert in the 3xTg-AD mouse model is significantly lower at 15 months of age [58]. The *hAPP*/*PSEN1*(ρEx9) mouse [59] and the 3xTg-AD mouse [60] suggest a loss of CB1R density is associated with progression of pathology. Given the ambiguities surrounding the role of the ECS in human and mouse health and disease, we chose to compare markers of the ECS in human autopsy AD brain samples to the same markers in a mouse model of AD-related amyloidosis. While we observed some similarities between human and mouse ECS in brain and how they tend to align with regards to distribution and expression, there were some clear differences. For example, we observed significant correlations between CB1R expression and MAGL in human cortical (but not hippocampal) samples. This co-distribution was evident whether samples were stratified by sex alone or by *APOE* ε4 status alone (sex [61] and *APOE* ε4 status [62, 63] being two major risk factors for AD-related pathology) and any co-distribution was lost when data were stratified by diagnosis. We did not observe the same association between FAAH and CB1Rs in our human brain samples. In mouse cortical and hippocampal samples, we observed co-distribution of CB1Rs with both MAGL and FAAH and these associations were not altered by the presence of the APP_Swe/Ind_ transgene. While such mouse models are often used to evaluate ECS function in an AD-like background, with quite some degree of success with respect to therapeutic outcomes, many of these outcomes do not translate efficiently to the clinical setting. Yet our observations do offer some options for consideration. For example, the fact that MAGL appears to align far better with levels of CB1R suggests that 2-AG (the substrate for MAGL) is more of an influence on CB1R in the human brain. The fact that this association is evident if data are simply stratified for sex or *APOE* ε4 status could establish criteria for personalized management options or inform on recruitment to clinical trials. We are currently evaluating how CB1R, FAAH, and MAGL align with levels of the AD-related Aβ peptide in our sample sets.

### Experimental procedures

#### Human brain samples

Autopsied cortical samples were obtained from the Douglas–Bell Canada Brain Bank (McGill University, Montréal, Canada) under the University of Saskatchewan’s Research Ethics Office Certificate of Approval: Bio 06-124 (DDM: Principal Investigator). Sixty male and female (M/F) samples matched as for age and sex include 26 controls (12M/14F) (see **Table 1**), 16 EOAD (*i.e.*, age of onset <65 years: 7M/9F), and 18 LOAD (*i.e.*, age of onset 65 + years: 8M/10F) (see **Table 2**). None of our cases of EOAD were carriers of any of the mutations in *PSEN1*, *PSEN2*, or *APP* genes that have been linked to the rare autosomal dominant forms of EOAD [64]. All cases of AD were confirmed neuropathologically at intake according to the CERAD criteria. Cortical samples corresponding to a mix of superior and middle frontal cortices (BA9/46, respectively) were chosen as they are associated with executive function and cognition, and relative hypoperfusion in AD patients [65]. Regional variation between markers of the ECS was explored by comparing our cortical samples with the corresponding hippocampal samples. Our hippocampal AD samples, *e.g.* EOAD (7M/8F), and LOAD (8M/10F), were well matched in number. Unfortunately, several of the control samples were depleted at source and we recognize this as a possible limitation of the study.

**Table 1:** Summary of neurologically normal, control donor information. De-identified information that was provided with the individual brain samples include the donor’s sex (M: male; F: female), Age (years), PMI (post-mortem interval, hours), COD (cause of death), Onset and Duration of disease (does not apply to control donors), *APOE* status (alleles ε2, ε3, or ε4), and weight (brain weight, grams). NI: no information.

| Sex | Age | PMI | COD | Onset | Duration | APOE | Weight |
| --- | --- | --- | --- | --- | --- | --- | --- |
| M | 89 | 22.75 | Heart failure |  |  | 3/3 | 935 |
| M | 70 | 32.75 | Hemorrhagic colitis |  |  | 3/4 | 1320 |
| M | 69 | 12 | Heart failure |  |  | 3/4 | 1225 |
| M | 69 | 27.5 | Heart failure |  |  | 2/4 | 1240 |
| M | 79 | 14.75 | Acute pulmonary edema |  |  | 3/3 | 1320 |
| M | 85 | 5.67 | Heart failure |  |  | 3/3 | 1080 |
| M | 60 | 7.25 | No discernable cause |  |  | 2/3 | 1350 |
| M | 61 | 8.75 | Pancreatic metastasis |  |  | 3/3 | 1337 |
| M | 59 | NI | Intestinal adenocarcinoma |  |  | 3/3 | 1443 |
| M | 64 | NI | Caecal carcinoma |  |  | 3/4 | 1530 |
| M | 77 | NI | Cerebral micro-infarcts* |  |  | 3/3 | 1255 |
| M | 66 | NI | Pancreatic adenocarcinoma |  |  | 3/3 | 1260 |
| F | 48 | 43.75 | Breast cancer metastasis |  |  | 3/4 | ND |
| F | 70 | 37 | Retroperitoneal hematoma |  |  | 3/3 | 1200 |
| F | 75 | 10.75 | Coronary arteriosclerosis |  |  | 3/3 | 1090 |
| F | 90 | NI | Cecal adenocarcinoma |  |  | 3/3 | 1050 |
| F | 77 | 7 | Pancreatic carcinoma |  |  | 3/3 | 1175 |
| F | 75 | 24.25 | Pulmonary edema |  |  | 3/3 | 1399 |
| F | 76 | 10.42 | Heart failure |  |  | 3/3 | 1260 |
| F | 82 | 32.58 | COPD |  |  | 3/3 | 1180 |
| F | 86 | 5.75 | Heart failure |  |  | 3/3 | 1192 |
| F | 50 | 11.5 | Heart failure |  |  | 3/3 | 1205 |
| F | 51 | 26.25 | Breast cancer |  |  | 3/3 | 1336 |
| F | 57 | 35.75 | Cecal adenocarcinoma |  |  | 3/3 | 1250 |
| F | 91 | NI | Pharyngeal neoplasm |  |  | 3/3 | 1000 |
| F | 63 | NI | Breast cancer |  |  | 3/3 | 1150 |

**Table 2:**
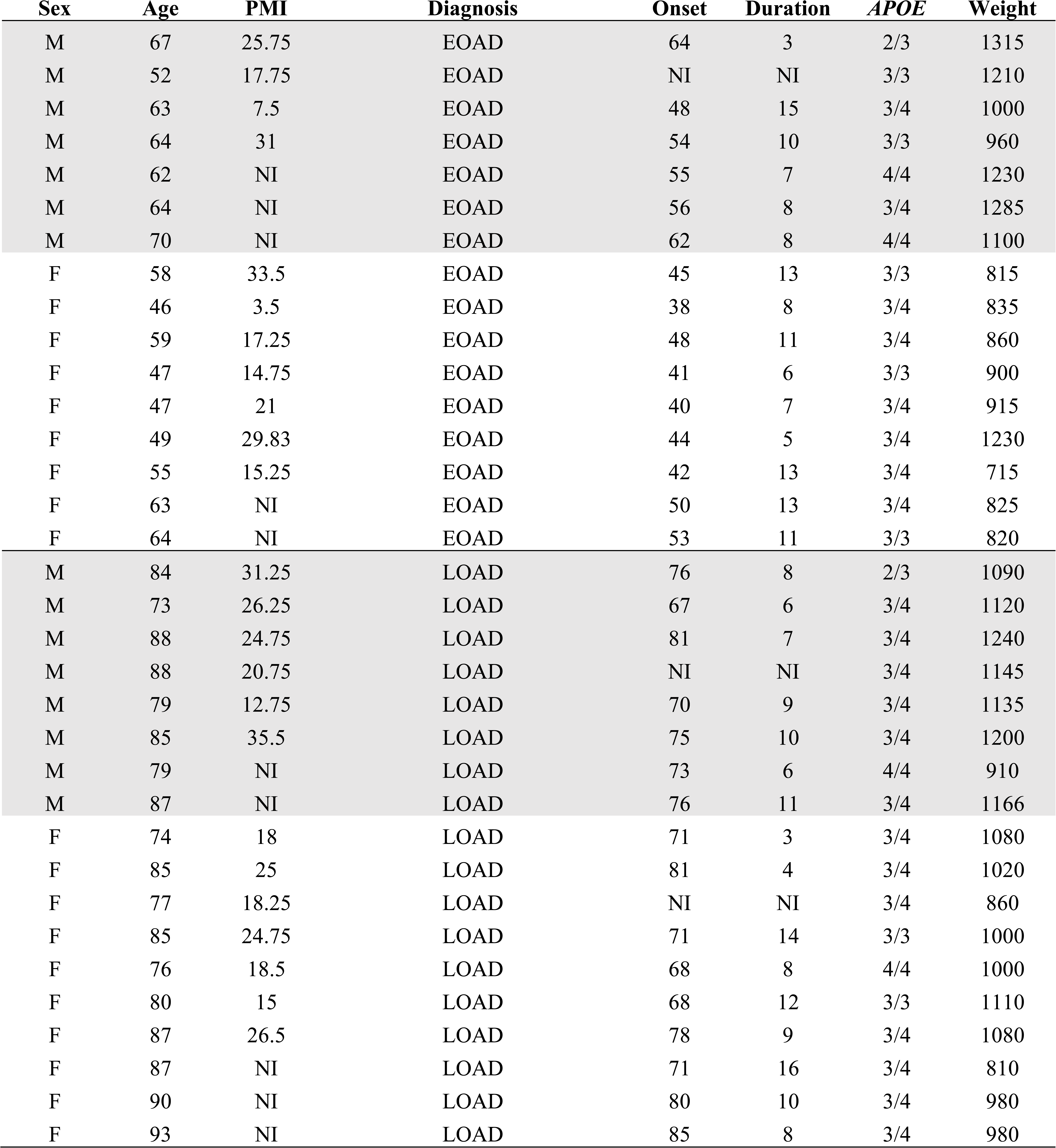
Summary of patient donor information. Donors included individuals with a diagnosis of early-onset AD (EOAD) or late-onset AD (LOAD), with the demarcation between classification as either EOAD or LOAD based on age of disease onset, *e.g.* EOAD = onset < 65 years; LOAD = onset > 65 years. De-identified information provided for each donor included: Sex (M: male; F: female), Age (years), PMI (post-mortem interval, hours), Onset of disease (age, years), Duration of disease (years), *APOE* status (alleles ε2, ε3, or ε4), and Weight (brain weight, grams). NI: no information.

#### APOE genotyping

We previously investigated the *APOE* ε4 status of each donor based on restriction isotyping [*e.g*., rs429358 (APOE-C112R) and rs7412 (APOE-R158C)] [63]. The donors’ *APOE* ε2/ε3/ε4 status is reported in **Tables 1 & 2**.

#### CNR1 (cannabinoid receptor 1) transcript expression using semi-quantitative PCR

Total RNA was isolated from pestle-homogenized brain tissue (50-100 mg) using the RNeasy® Plus Universal Mini Kit (Qiagen; cat#: 73404). RNA (1 μg: quantified using nano-drop spectrophotometry) was reverse-transcribed using the iScript Select cDNA Synthesis Kit (Bio-Rad; cat# 170-8897) and cDNA was quantified by nanodrop spectrophotometry. Transcripts of interest were amplified by standard polymerase chain reaction (PCR) using 500 ng of the respective cDNA and Platinum Taq DNA Polymerase (Invitrogen; cat#10966-034). The Forward (F) and Reverse (R) primer pairs for the coding sequence were based on GenBank sequence (accession# BC100968.1) as reported elsewhere [66], whereas the primer pair spanning Exon1-Exon4 were designed based on human *CNR1*, transcript variant 1, NCBI (NM_016083.4). The primer pairs were: (a) *CNR1* (amplicon = 165 bp): (F) 5’-AAG GTG ACA TGG CAT CCA AAT and (R) 5’-AGG ACG AGA GAG ACT TGT TGT AA; (b) *CNR1* Spanning Intron (amplicon = 265 bp): (F) (in Exon 1) 5’-AGG ACC AGG GGA TGC GAA GG and (R) (in Exon 4) 5’-AAT TTG GAT GCC ATG TCA CCT T; (c) Succinate dehydrogenase (SDH: used as internal control [67]; amplicon = 141 bp) (F) 5’-TCT GCC CAC ACC AGC ACT and (R) 5’-CCT CTC CAC GAC ATC CTT CC. Thermocycling relied on a denaturing step at 95°C (10 minutes), followed by 50 cycles of 95°C/58°C/72°C (10 seconds at each temperature), with a final extension at 72°C for 5 minutes. 10 μL of the reaction mixture was then resolved on a 2% agarose gel containing 5 μL/100mL GelRed Stain (Cat# 41003-Biotium) and run at 80V for 1 Hr (100 mL gel) in 0.5X TBE. Amplicons were visualized under UV light and quantified by densitometry (ImageJ 1.32j: http://rsb.info.nih.gov/ij/).

#### Antibodies and immunodetection

The anti-CB1R antibody (Sigma-Aldrich: cat#C1108) and anti-FAAH (WH0002166M7) antibodies were obtained from Sigma-Aldrich Ltd, while anti-MAGL (Ab24701) was obtained from Abcam Inc. The GAPDH antibody (#14C10) was obtained from Cell Signaling Technologies. LICOR goat anti-rabbit or anti-mouse secondary antibodies were obtained from Bio-Rad Laboratories (Canada) Ltd.

The RIPA buffer used for all autopsied sample preparations contained 50 mM Tris (pH7.5), 150 mM NaCl, 1% Nonidet P-40 (NP-40), 0.5% Sodium Deoxycholate, 1 mM EDTA, 0.1% SDS with protease and phosphatase inhibitors (added fresh) 1 mM PMSF, 2 mM Na_3_VO_4_, 5 mM NaF, 5 mM Sodium Pyrophosphate (Sigma-Aldrich Ltd, 4693132001). Immunodetection conditions were first optimized using different denaturing temperatures and combinations of milk casein or BSA blocking buffers and/or milk casein or BSA buffer for incubation with the primary antibodies. In our hands, the signal-to-noise was far better when using BSA as a blocking agent.

Tissue samples (∼30 mg) were disrupted in 20 volumes of RIPA using a pestle-homogenizer system and then passed five times through a 22-gauge needle. Samples were allowed to sit on ice for 30 minutes and then precleared by centrifugation (12,000×*g*, 10 min, 4 °C) and used for protein determination (Sigma Total Protein Kit: cat#TP0300). Samples were diluted in Laemmli Buffer to 1 μg/μl and heated for five minutes at 65°C (the lower temperature still denatures proteins, but avoids aggregation of glycosylated proteins, *e.g.* the CB1R): [68]). This assay was performed by a single individual so as to decrease inter-sampling variability.

Sample aliquots (25 μg protein/lane) were resolved by SDS-PAGE and then transferred to nitrocellulose membranes (230 mA, 90 minutes) and blocked with 1% BSA in TBST (TBS +0.2% Triton X-100) for 30 minutes. Membranes were then washed in TBST (3×10 minutes) and probed overnight at 4°C with primary antibody [anti-CB1R at 1:500; anti-FAAH at 1:1000; anti-MAGL at 1:1000]. Membranes were then washed in TBST and incubated with LICOR goat anti-rabbit-800 secondary (1:20,000 in 5% milk/TBST) for one hour at room temperature. Membranes washed in TBST (3×10min) and scanned on an Odyssey Imager (LI-COR Biosciences). The expression of total GAPDH was used to monitor protein loading across samples.

#### The ‘J20’ APP mouse model of AD-related amyloidosis

The B6.Cg-Tg(PDGFB-*APPSwInd*)20Lms/2Mmjax (J20) mouse strain (#06293: Jackson Laboratory) expresses a human *APP695* gene variant coding for the Swedish (K670N/M671L) and Indiana (V717F) substitutions [69]. We have detected the β-amyloid (Aβ) peptide in male and female J20 mice as young as 3 months of age [70]. Animal tissues –based on 15-month-old heterozygous J20 mice and their wildtype (WT) littermates– were collected with approval by the University of Saskatchewan’s Animal Research Ethics Board (protocols 20040068 & 20060070: PI - DDM). Frontal cortex and corresponding hippocampal lysates were analyzed for CB1R, FAAH, and MAGL protein expression using standard SDS-PAGE immunoblotting (as above). Tubulin-III (Sigma-Aldrich cat#T8578) was used as a loading control.

### Statistical analyses

The variability between group means was analyzed using either the Mann-Whitney U test or two-way ANOVA (α = 0.05) with *post hoc* multiple comparisons based on the Tukey test. In addition to comparing sample means based on diagnosis, *e.g.* control, EOAD, LOAD, we also examined the data based on stratification into homogeneous subgroups independent of diagnosis, *e.g.* by sex alone or by *APOE* ε4 status alone. Simple linear regression was used to estimate any association between test variables. Significance was set at *P*<0.05; however, values between 0.051 and 0.099 were considered *tendencies*. Due to the volume of data, most summaries of any analyses based on stratification that did not result in any overt statistically relevant patterns are supplied as *Supplementary Tables* (see **Supp Tables 1-27**). Any statistically significant effects are discussed in the text and in the corresponding figure legend. Note that our sample set allowed us to analyze the data by diagnosis alone, sex alone, *APOE* ε4 status alone (carrier or not of the ε4 risk allele), sex-by-diagnosis, or sex-by-*APOE* ε4 status. Unfortunately, it did not allow us to stratify the data based on diagnosis-by-sex-by-*APOE* ε4 status, *e.g*. our random sample set only included a single *APOE* ε4(+) female control sample and a single *APOE* ε4(-) male LOAD sample (see **Tables 1 and 2**; donor statistics).

## Results

### CNR1 transcripts in human cortical extracts

Semi-quantitative PCR was used to estimate mRNA transcript levels of *CNR1* (the gene encoding the CB1R) in human brain (**Figure 1**). Two sets of primer-pairs led to distinct patterns. Transcript levels of succinate dehydrogenase (SDH, *e.g.* Complex II) were used to normalize amplicon intensity. A primer-pair within *CNR1* Exon 4, which contains the CB1R protein coding region, and is the primary exon expressed in brain [71], revealed a fairly consistent pattern of expression across samples and no observable differences between samples means whether stratified by diagnosis, by *APOE* ε4 status or by sex alone (**Figure 1B-D**). In these same samples, we investigated the pattern of expression based on a primer-pair covering Exon 1 and Exon 4. We chose to examine this combination as Exon 1 contains multiple transcription starting sites [72] that may contribute to alternative *CNR1* transcription and CB1R function [73]. Exon 1-4 amplicons were missing from some samples (suggesting a lack of Exon 1 transcription), whereas other samples produced slightly different amplicon sizes (**Figure 1A**).

**Figure 1.**
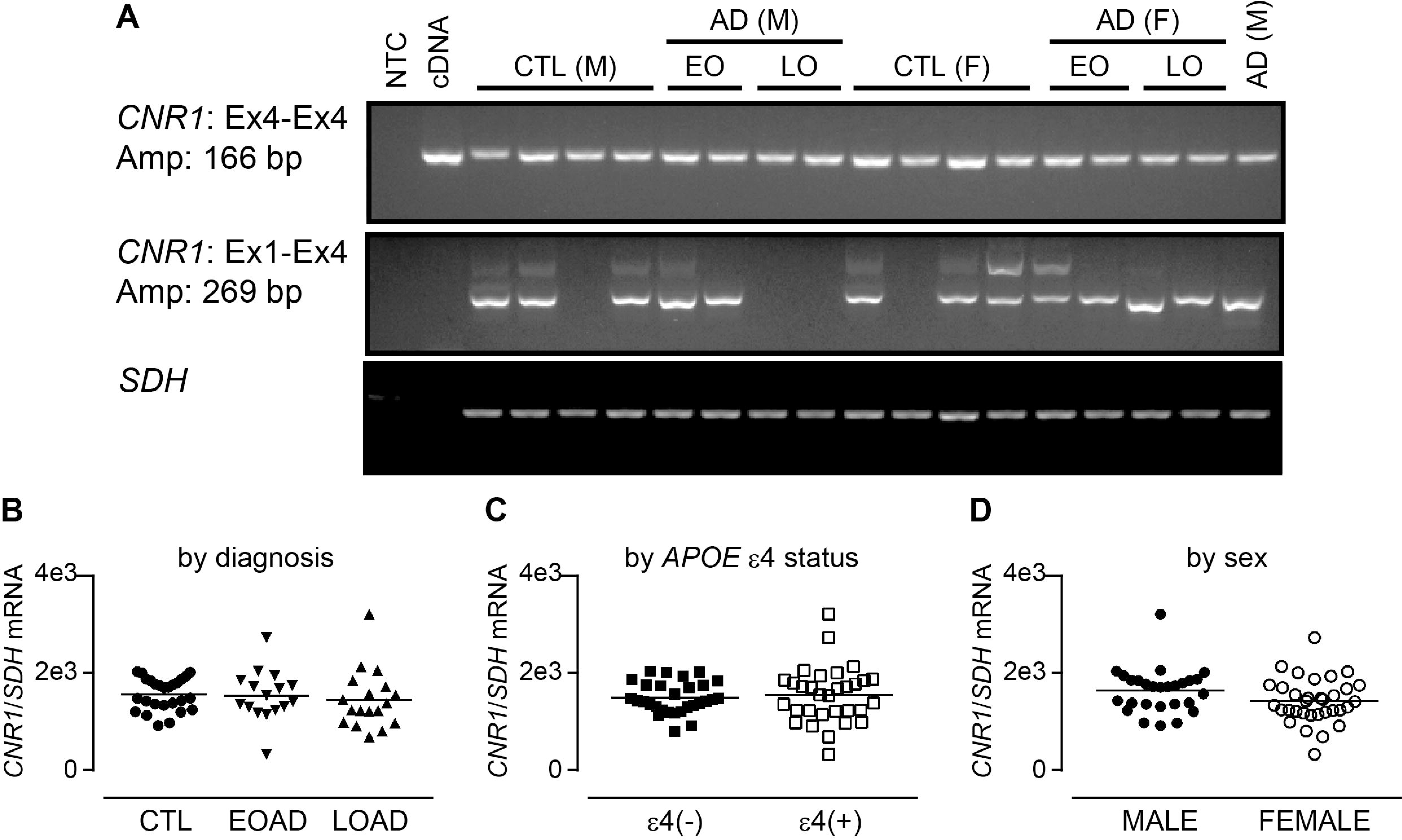
CNR1 transcripts in human control and Alzheimer disease brain samples. (**A**) Amplicons generted primer pairs within Exon 4 (Ex4) and spanning Exon1 and Exon4 of CNR1 cDNA in neurologically normal control (CTL), early-onset AD (EO), and late-onset AD (LO) male (M) and female (F) samples. Levels of a succinate dehydrogenase (*SDH*) amplicon was used as an internal control. NTC (no template control) and cDNA (a plasmid construct containing the *CNR1* Exon4 cDNA) were used as negative and positive controls, respectively. The relative expression of *CNR1*(Ex4) to *SDH* amplicons are expressed (**B**) by the donor’s diagnosis, (**C**) by the donor’s *APOE* ε4 status, *e.g.*, non-carriers: ε4(-); carriers: ε4(+), or (**D**) by the donor’s sex alone.

### Western blot analysis of components of the ECS in human brain

#### Comparing data based on group means

Many phenotypes associated with the ECS across various neurodegenerative disease, including AD, have been based on mRNA transcript levels, receptor binding and pharmacological approaches, with a certain degree of ambiguity between studies (reviewed in [4]). We chose to investigate the expression of the CB1R protein in our human autopsy sample set (cortex and hippocampus). Western blotting revealed a pattern that included three anti-CB1R immunoreactive bands (**Figure 2A**). Based on published reports, the 47-kDa form represents the immature CB1R (*e.g*., with no post-translational modifications), whereas the 60-kDa form likely represents a glycosylated (putatively active) form [74]. We also detected a triplet that migrated at approximately 37-kDa band; this band grouping has been described elsewhere as likely representing a putative splice variant rather than any degradation fragment(s) [75]. Aside from an increase in the 60-kDa CB1R band (CB1R-60) in the hippocampus of our EOAD and LOAD samples [P = 0.0088] (**Figure 2B**), we did not observe any changes in the mean expression of any of these CB1R bands between controls or other AD samples (**Figure 2B-D**). ANOVA did not reveal any overt changes in mean expression levels of either cortical or hippocampal FAAH (**Figure 2E**) or MAGL (**Figure 2F**), regardless of diagnosis. This lack of central tendencies based on group means reflects the reports of inconsistent changes in markers of the ECS in AD tissues [4]. We chose to use linear regression to question whether any variability around the means could be indicating associations in the expression of the different markers of the ECS.

**Figure 2.**
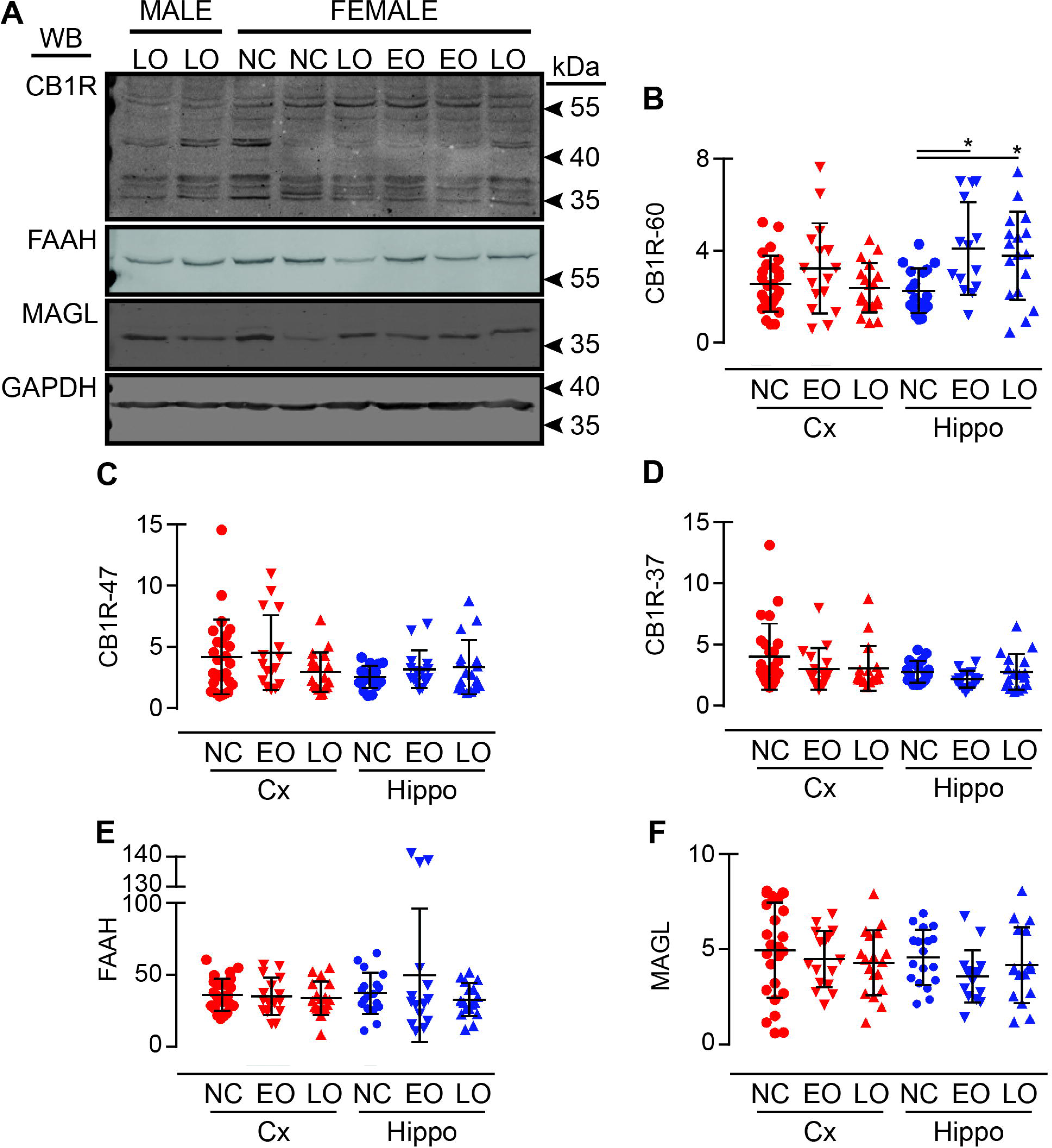
Expression levels of markers of the ECS in human cortical and hippocampal samples stratified by diagnosis. (**A**) Representative immunoblots of Cannabinoid Receptor 1 (CB1R), FAAH, MAGL, and GAPDH (loading control) in human neurologically normal control (NC), early-onset AD (EO), and late-onset AD (LO) samples. The anti-CB1R antibody reveals three CB1R immunodetetctable bands at ∼60 kDa, ∼47 kDa, and ∼37 kDa. The levels (*y*-axis expressed as ‘relative units’) of (**B**) CB1R-60, (**C**) CB1R-47, (**D**) CB1R-37, (**E**) FAAH, and (**F**) MAGL in cortical (Cx) and hippocampal (Hippo) samples are presented by diagnosis. *: *P* < 0.05 between indicated groups.

### Using linear regression to determine whether associations exist between the components of the ECS in human brain

#### Analyses based on data stratified by diagnosis

As stated in the ‘*Statistics*’ section, we chose to relegate analyses without any overt statistical significance to supplementary tables to limit the number of figures in the main body of the text. The term ‘pooled data’ refers to the pooling of male and female data. Somewhat unexpectedly, our regression analyses ‒when data were stratified for diagnosis‒ revealed that any associations between the expression of any of the CB1R species were sporadic and were not confined to control or AD samples. For example, we observed a significant association between cortical CB1R-60 and CB1R-37 (P = 0.032) and CB1R47 and CB1R-37 (P = 0.034) in control samples and these were driven by females in both cases (P = 0.008 and P < 0.000, respectively; **SUPPL Table 1**). None of these associations were observed in the hippocampal samples (**SUPPL Table 2**). When comparing CB1R expression levels between regions, we only observed a strong association between cortical and hippocampal CB1R-37 expression in EOAD samples (pooled: P = 0.003; male – P = 0.564; females – P = 0.003) (**SUPPL Table 3**).

#### Alignment of different CB1R bands within regions based on data stratified by APOE ε4 status (e.g. carrier or not)

At this point, given that some of our observations indicated clear sex differences in patterns, we chose to exclude ‘diagnosis’ as a nominal variable. We first compared group means when CB1R data were stratified simply by whether the donor was a carrier of the *APOE* ε4 risk allele for AD or by sex alone. Except for a significant increase in CB1R-60 expression in hippocampal samples of carriers of the ε4 allele [P = 0.0044], the expression levels of CB1Rs were effectively unchanged in our cortical and hippocampal samples (**SUPPL Figure 1**). Similarly, we did not observe any effect of sex alone on the expression of CB1Rs (**SUPPL Figure 2**). We did not observe any significant changes in expression of CB1Rs when cortical data were stratified by *APOE* ε4-by-sex (**Figure 3A-C**), whereas the significant increase we observed in hippocampal CB1R-60 in carriers of the ε4 allele (**SUPPL Figure 1**, see above) was influenced primarily by female carriers of the allele (male +ε4: P = 0.1348; female +ε4: P = 0.0120] (**SUPPL Figure 3).**

**Figure 3.**
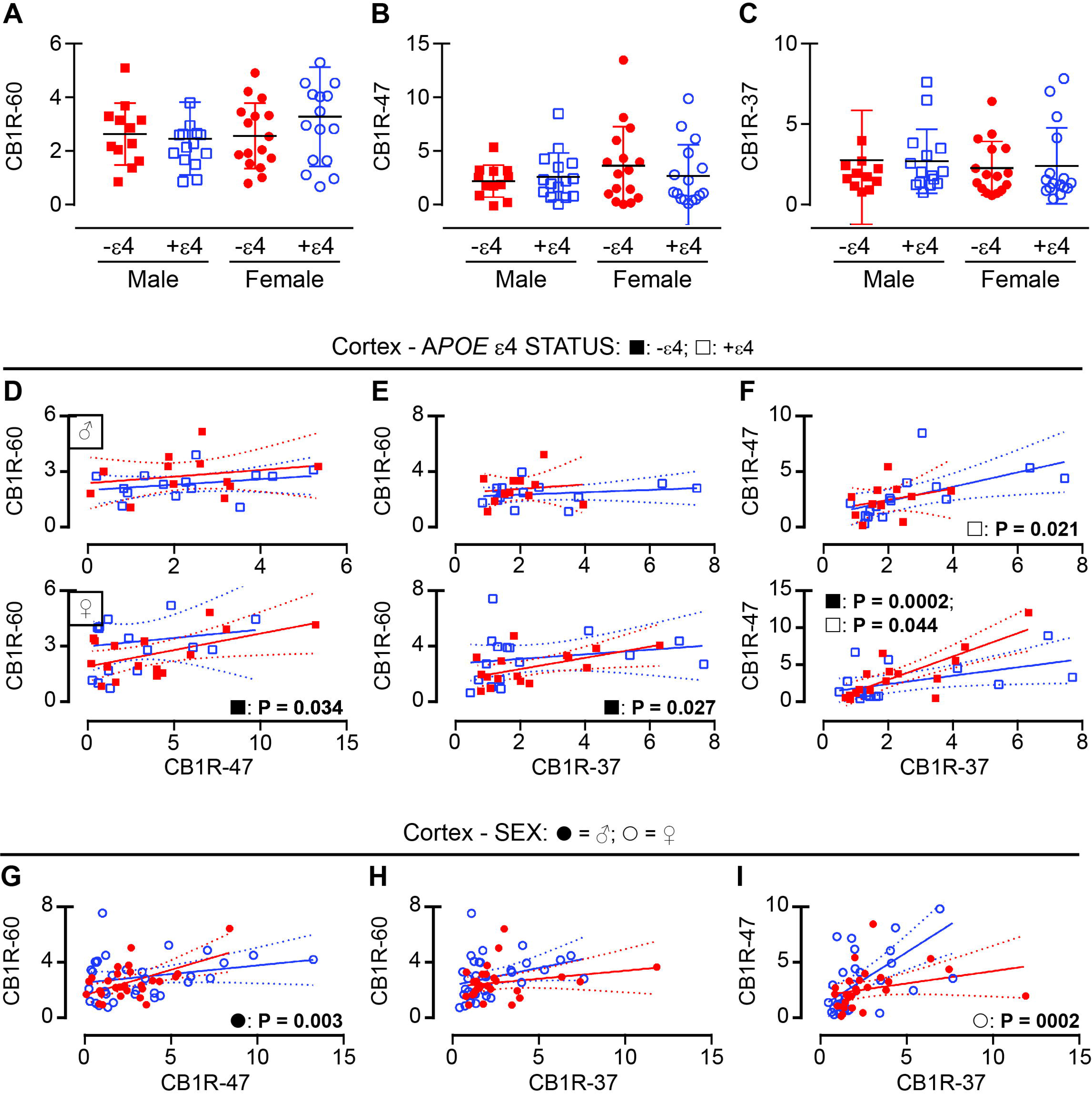
Expression levels of CB1Rs stratified by *APOE* ε4 status or by sex alone. The levels of (**A**) CB1R-60, (**B**) CB1R-47, and (**C**) CB1R-37 in cortical samples grouped according to the donor’s *APOE* ε4 status, *e.g.*, non-carriers: ε4(-) [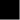]; carriers: ε4(+) [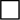]. The same data were used for regression analysis between (**D**) CB1R-60 and CB1R-47, (**E**) CB1R-60 and CB1R-37, and (**F**) CB1R-47 and CB1R-37, with males depicted in the top row and females in the bottom row. The association (liner regression) between (**G**) CB1R-60 and CB1R-47, (**H**) CB1R-60 and CB1R-37, and (**I**) CB1R-47 and CB1R-37 when data are stratified by sex alone [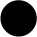: male; ○: female]. Only those associations that are significant (*P* ≤ 0.05) are identified (next to their grouping symbol).

We observed a strong positive association between CB1R-60 and CB1R-47 (P = 0.034) and between CB1R-60 and CBR1-R37 (P = 0.0268) solely in the cortex of female non-carriers of the ε4 allele (**Figure 3D, E**). Unexpectedly, we also observed an association between CB1R-47 and CB1R-37 in male carriers (ε4 neg: P = 0.3727; ε4 pos: P = 0.0210) and in females regardless of allele status (ε4 neg: P = 0.0002; ε4 pos: P = 0.0442) (**Figure 3F**). In contrast, we only observed an association (in this case a negative association) between CB1R-60 and CB1R-37 expression in hippocampal samples and this was limited to male non-carriers of the allele (P = 0.0179) (**SUPPL Table 8**).

#### Alignment of different CB1R bands within regions based on data stratified by sex alone

If analyzed by sex alone (*e.g*., independent of *APOE* ε4 status or diagnosis), we observed a strong association between CB1R-60 and CB1R-47 in males (P = 0.0029) (**Figure 3G**) and between CB1R-47 and CB1R-37 in females (P = 0.0002) (**Figure 3I**). We did not observe any association between CB1R-60 and CB1R-37 in cortex (**Figure 3H**), nor between any of the CB1R species in the hippocampus (**SUPPL Table 5**). The only association between cortical and hippocampal CB1R species when data were stratified by *APOE* ε4 status and/or sex was the expression of CB1R-37 in females, which was maintained whether the data were stratified by *APOE* ε4 status (male ε4 pos: P = 0.576; female ε4 pos: P = 0.008) or by sex alone (male: P = 0.367; female: P = 0.025) (**SUPPL Table 6**).

There is clear evidence in the literature that inhibition of components of the ECS can impact CB1R function, presumably through a change in endogenous ligand availability and subsequent receptor down-regulation [55, 56]. As such, we included FAAH and MAGL in our screen of markers of the ECS. As stated above, we did not detect any overt changes in mean expression levels of either FAAH (**Figure 2E**) or MAGL (**Figure 2F**) in either region, regardless of diagnosis.

#### Alignment of FAAH and MAGL within regions based on data stratified by diagnosis

There was no overt associations between FAAH and MAGL levels when data were stratified by diagnosis, e.g. control, EOAD, or LOAD, with the exception of control males in hippocampus [P = 0.035] (**SUPPL Table 9**). We also did not observe any association between cortical and hippocampal levels of either protein when stratified by diagnosis, with the exception of pooled control samples [P = 0.029] (**SUPPL Table 10**).

#### Alignment of FAAH and MAGL within regions based on data stratified by APOE ε4 status (e.g. carrier or not)

Focussing exclusively on FAAH and MAGL expression levels when data were grouped by *APOE* ε4 status alone (**SUPPL Figure 4A, B**) or sex alone (**SUPPL Figure 4C, D**) did not reveal any difference between carriers and non-carriers in either cortical or hippocampal FAAH levels. This lack of any significant difference in expression levels of either protein extended to stratification by *APOE* ε4-by-sex (**Figure 4A, B**), although ANOVA did reveal an interaction [P = 0.0366] between *APOE* ε4 status and sex in our cortical MAGL samples (**Figure 4A**), but not in the corresponding hippocampal samples (**Figure 4B**). We observed no association between the expression of the two enzymes in cortex if stratified by *APOE* ε4-by-sex (**Figure 4C, D**) and only a modest association between the two enzymes in females, if stratified by sex alone (P = 0.0494) (**Figure 4E**). We did observe a modest association between FAAH and MAGL expression in hippocampal samples from male non-carriers of the allele (P = 0.0318) and female carriers of the allele (P = 0.0421) (**Figure 4F, G**), but this was lost when stratified by sex alone (**Figure 4H**). There were no observed association between cortical or hippocampal FAAH (or cortical or hippocampal MAGL), whether stratified by *APOE* ε4-by-sex or by sex alone (**SUPPL Table 11**).

**Figure 4.**
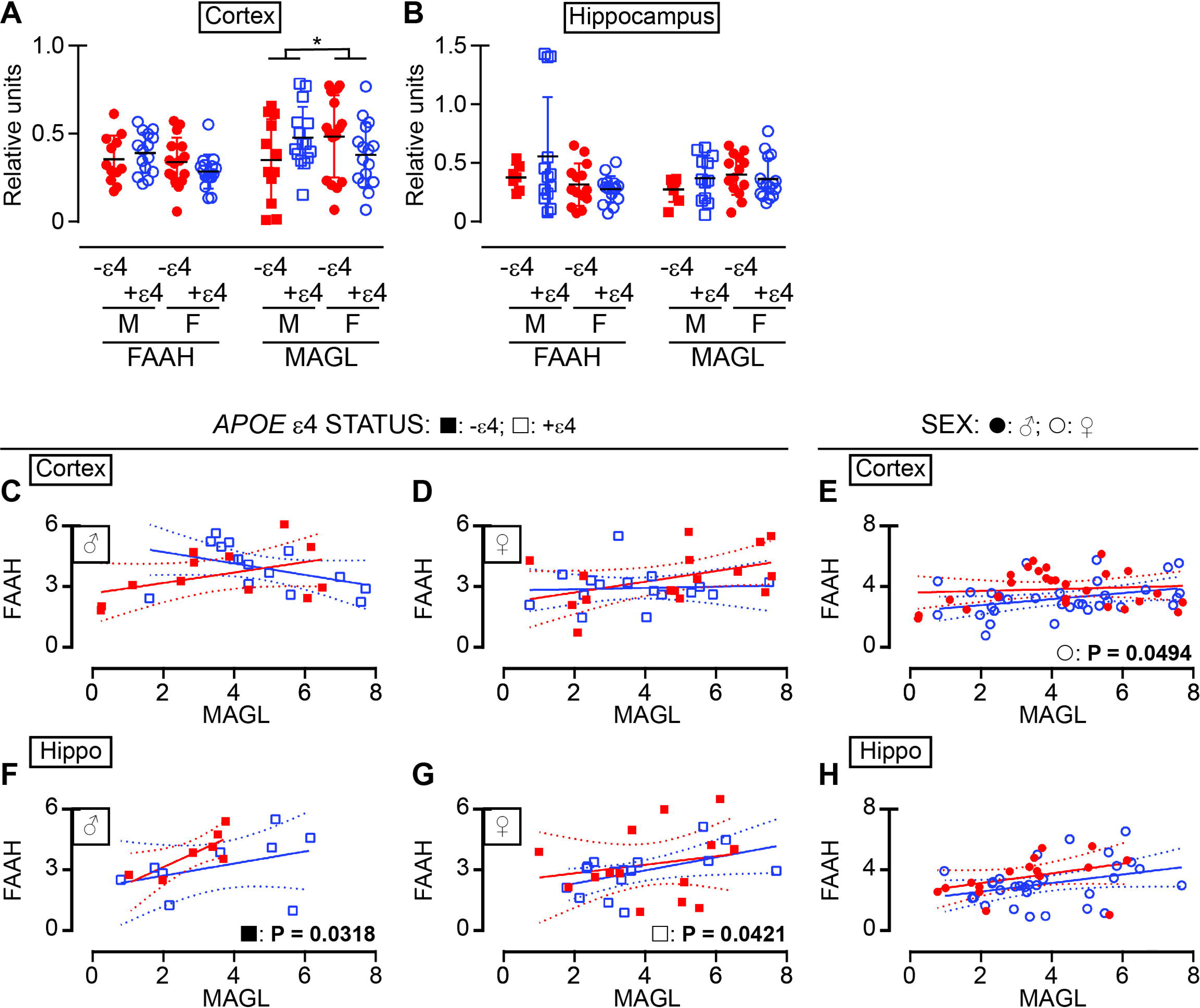
Expression levels of FAAH and MAGL stratified by *APOE* ε4 status or by sex alone in cortical and hippocampal samples. The levels of (**A**) FAAH and MAGL in cortical samples grouped according to the donor’s *APOE* ε4 status, *e.g.*, non-carriers: ε4(-); carriers: ε4(+), and sex. The levels of (**B**) FAAH and MAGL in corresponding hippocampal samples. The association (liner regression) between FAAH and MAGL in male and female (**C**, **D**) cortical and (**F**, **G**) hippocampal samples stratified by *APOE* ε4 status, *e.g.*, non-carriers: ε4(-) [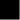]; carriers: ε4(+) [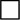]. (**E**, **H**) The same datapoints were used for regression when stratified by sex alone [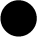: male; ○: female]. *: interaction between *APOE* ε4 status and sex (MAGL in cortex). Only those associations that are significant (*P* ≤ 0.05) are identified (next to their grouping symbol).

#### Alignment of FAAH and MAGL within regions based on data stratified by sex alone

Aside from a tendency for an association in male cortical samples [P = 0.077], we did not observe any associations between the proteins when stratified for sex alone (**SUPPL Table 12**).

We next chose to investigate whether the expression of CB1R immunodetectable bands aligned with the expression of either FAAH or MAGL using our stratifications, *e.g*. diagnosis, *APOE* ε4 status, or sex.

#### Alignment of either FAAH or MAGL and CB1Rs within regions based on data stratified by diagnosis

Linear regression revealed sporadic associations between cortical CB1R-47 and FAAH in female LOAD samples (P = 0.011; **SUPPL Table 13**) and between hippocampal CB1R-47 and FAAH in control (pooled) samples (P = 0.022; **SUPPL Table 14**). Interestingly, we did observe more frequent associations between MAGL and CB1Rs. For example, CB1R-47 associated with MAGL in cortex (control males: P = 0.002; EOAD females: P = 0.015) (**SUPPL Table 15**) and with MAGL in hippocampus (control males: P = 0.005; control females: P = 0.027; EOAD females: P = 0.032; LOAD males: P = 0.025) (**SUPPL Table 16**).

Finally, given the lack of any generalized patterns of association between CB1Rs and FAAH or MAGL based on a diagnosis of EOAD or LOAD, we investigated whether levels of FAAH (or MAGL) associated with expression of the CB1R based on the donors *APOE* ε4 status or their sex (independent of any diagnosis).

#### Alignment of FAAH and CB1Rs within regions based on data stratified by APOE ε4 status (e.g. carrier or not) or sex alone

We did not observe any association between FAAH expression and any of the CB1R immunoreactive bands in the cortex when stratified by *APOE* ε4 status (**SUPPL Table 17**) and only a weak association between FAAH and CB1R-47 in females (P = 0.0499) when analyzing by sex alone (**SUPPL Table 18**). We observed a similar lack of association between FAAH and CB1Rs in hippocampal tissues when stratified by *APOE* ε4 status (**SUPPL Table 19**) and a tendency for an association between FAAH and CB1R-47 in females (P = 0.0759) when analyzing by sex alone (**SUPPL Table 20**).

#### Alignment of MAGL and CB1Rs within regions based on data stratified by APOE ε4 status (e.g. carrier or not) or sex alone

In contrast to the generalized lack of association between FAAH and CB1R species reported above, we observed an association between MAGL levels and CB1R47 levels in non-carriers of the *APOE* ε4 allele (males: P 0.0228; females: P = 0.0548) (**Figure 5B**) and between MAGL and CB1R-37 in carriers of the ε4 allele (males: P = 0.0027; females: P = 0.0031) (**Figure 5C**). Levels of CB1R-60 did not align with the expression of MAGL in these samples (**Figure 5A**). Levels of MAGL were also very strongly (and selectively) associated with levels of CB1R-47 in both male (P = 0.0142) and female P = 0.0064) cortical samples when stratified by sex alone (**Figure 5E**), whereas hippocampal samples only exhibited a strong association between CB1R-47 in male carriers of the ε4 allele (P = 0.0019) (**SUPPL Table 21**) or only in males (P = 0.0009) when sex alone was considered (**SUPPL Table 22**).

**Figure 5.**
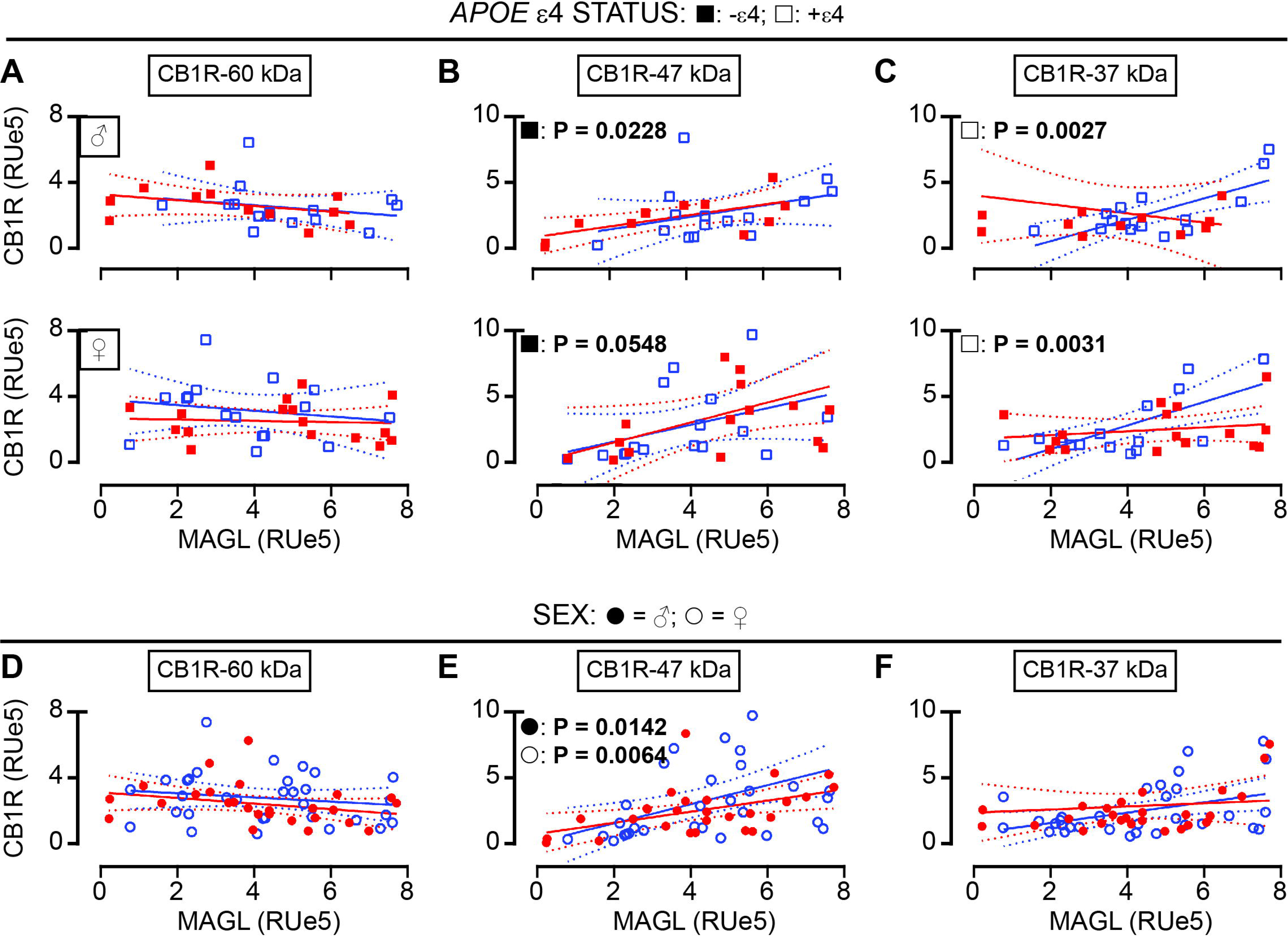
Association between CB1Rs and MAGL levels stratified by *APOE* ε4 status or by sex alone in cortical samples. The association (linear regression) between (**A**) CB1R-60, (B) CB1R-47, and (C) CB1R-37 and MAGL in cortical samples stratified by the donor’s *APOE* ε4 status, *e.g e.g.*, non-carriers: ε4(-) [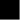]; carriers: ε4(+) [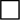]. (**D**, **E**, **F**) The same datapoints when stratified by sex alone [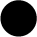: male; ○: female]. Only those associations that are significant (*P* ≤ 0.05) are highlighted (next to their grouping symbol).

Our dataset suggested that any association between markers of the ECS, such as CB1Rs, FAAH, and MAGL, in human brain tended to be influenced by the donor’s *APOE* ε4 status, and more often, simply based on the donor’s sex, and any association tended to be most obvious in cortex. Importantly, most associations were disrupted when data were stratified by diagnosis, whether it be EOAD or LOAD. It was also quite evident that levels of CB1Rs tend to associate more so with levels of MAGL (rather than FAAH), which suggests *a priori* that these receptors respond much more readily to the functionality of MAGL (and its primary substrate 2-AG).

Given the routine use of mouse models to investigate the contribution of the ECS to AD-related pathology, we chose to compare the patterns of ECS marker expression observed using human samples to the same markers in cortical and hippocampal tissues from male and female ‘J20’ mice (a model of AD based on the hAPP_Swe/Ind_ transgene [69]) and their wildtype (WT) littermates.

### Western blot analysis of ECS components in mouse brain

#### Comparing mouse data based on group means

We include an example of an immunoblot depicting the expression of markers of the ECS in the mouse brain (**Figure 6A**). Two-way ANOVA did not reveal any change in the expression of CB1R-60 in either the cortex or corresponding hippocampal samples (**Figure 6B**), regardless of the mouse genotype (*e.g.* wildtype or J20/transgenic). However, in clear contrast with our human data above, we observed a strong influence of sex on hippocampal CB1R-47 expression [P = 0.0017] as well as an interaction between sex and genotype in this same sample set [P = 0.0019] (**Figure 6C**). We also observed strong influence of sex on mean cortical [P = 0.0003] and hippocampal [P < 0.0001] CB1R-37 expression (and a strong *interaction* between sex and genotype in the latter set [P = 0.0001] (**Figure 6D**). There was also a strong influence of sex on mean cortical [P < 0.0001], but not hippocampal, FAAH expression (**Figure 6E**) as well as effects of sex on mean cortical [P = 0.0306] and hippocampal [P = 0.0133] MAGL expression (**Figure 6F**).

**Figure 6.**
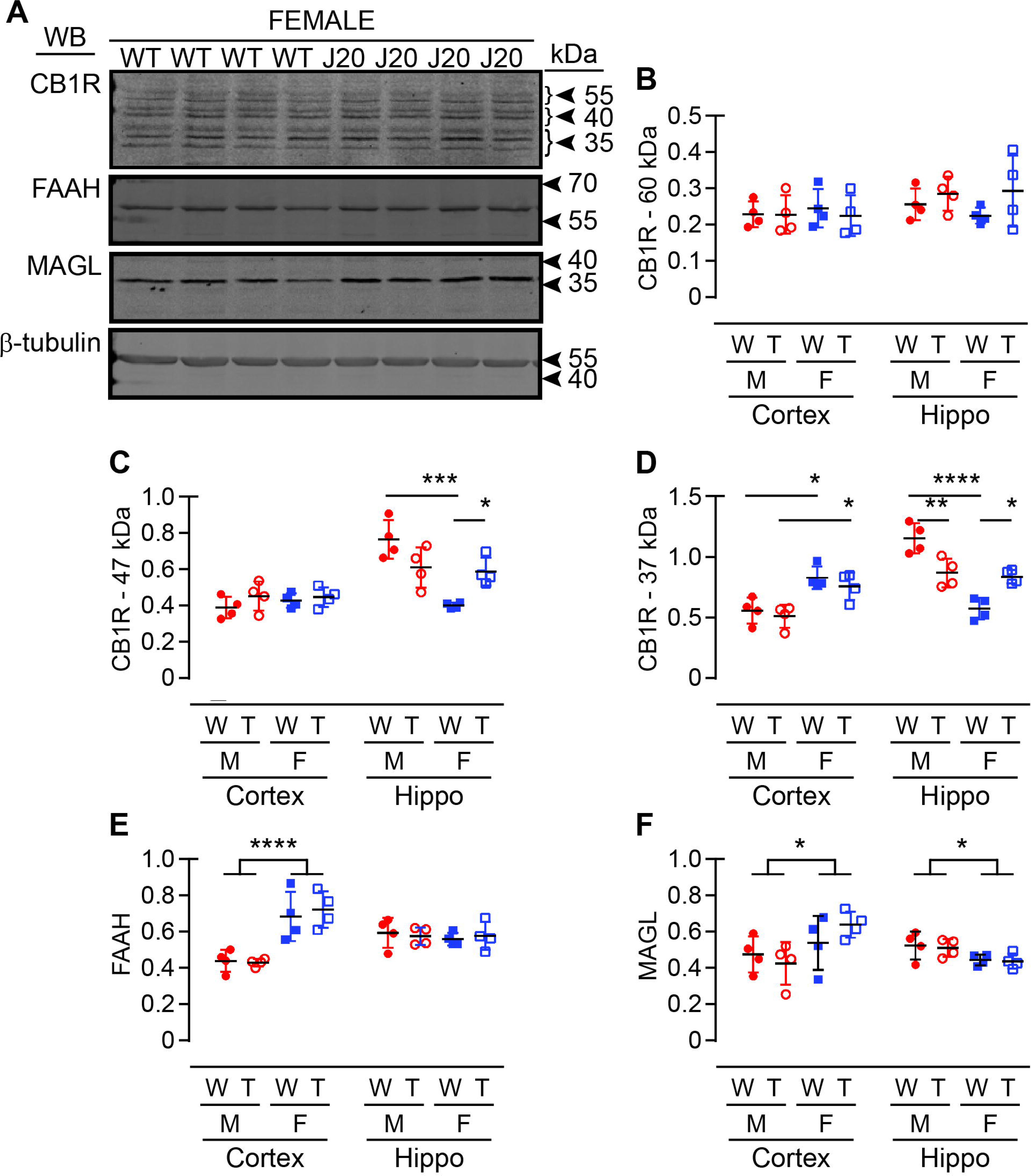
Expression levels of markers of the ECS in wildtype and J20 mouse brain. (**A**) Representative immunoblots of CB1R, FAAH, MAGL, and β-Tubulin (loading control) in mouse wildtype (WT) and hAPP_Swe/Ind_ (‘J20’) mice. The anti-CB1R antibody reveals a similar pattern of three CB1R immunodetetctable bands at ∼60 kDa, ∼47 kDa, and ∼37 kDa as seen in the human samples (see Figure 2). The levels (*y*-axis expressed as ‘relative units’) of (**B**) CB1R-60, (**C**) CB1R-47, (**D**) CB1R-37, (**E**) FAAH, and (**F**) MAGL in cortical and corresponding hippocampal samples are presented by gentype. *: *P* < 0.05; **: *P* < 0.01; ***: *P* < 0.001; ****: *P* < 0.0001 between indicated groups

#### Using linear regression to determine associations between expression of components of the ECS in mouse brain

*Alignment of mouse CB1R-60/-47/-37 levels based on pooled data (regardless of sex or genotype) or by stratification by genotype (e.g. WT or the J20 carrier of the hAPP_Swe/Ind_ transgene) and/or by sex*. The only association we observed between regions was the expression of CB1R-37 when all data were pooled (regardless of sex or genotype) [P = 0.006] or in males when data were stratified by sex alone [P = 0.0117] **(SUPPL Table 23)**. Within the cortex itself, CB1R-47 associated with CB1R-37 in WT mice [P = 0.0348] (**SUPPL Table 24**), while in the hippocampus, we observed an association between CB1R-60 and CB1R-47 in female mice [P = 0.0314] and strong associations between CB1R-47 and CB1R-37 whether all data points were pooled [P < 0.0001], or stratified either by sex alone [male: P = 0.0030; female: P = 0.0018] or by genotype [WT: P < 0.003; J20: P = 0.002] (**SUPPL Table 25**).

#### Alignment of mouse FAAH or MAGL levels based on stratification by genotype and/or by sex

There was no association between cortical and hippocampal FAAH levels when data were pooled or stratified by sex or genotype alone (**Figures 7A-C**), although breaking down the ‘genotype’ by sex revealed a strong association limited to the male J20 mice [male: P = 0.0107; female: P = 0.6087] (included in **Figure 7C**). We also observed an association between cortical and hippocampal MAGL levels if the data were pooled (regardless of sex or genotype) [P = 0.0397] (**Figure 7D**), but not when stratified by sex alone (**Figure 7E**). We did observe an association between cortical and hippocampal MAGL levels in J20 mice, regardless of sex [P = 0.0040] (**Figure 7F**).

**Figure 7.**
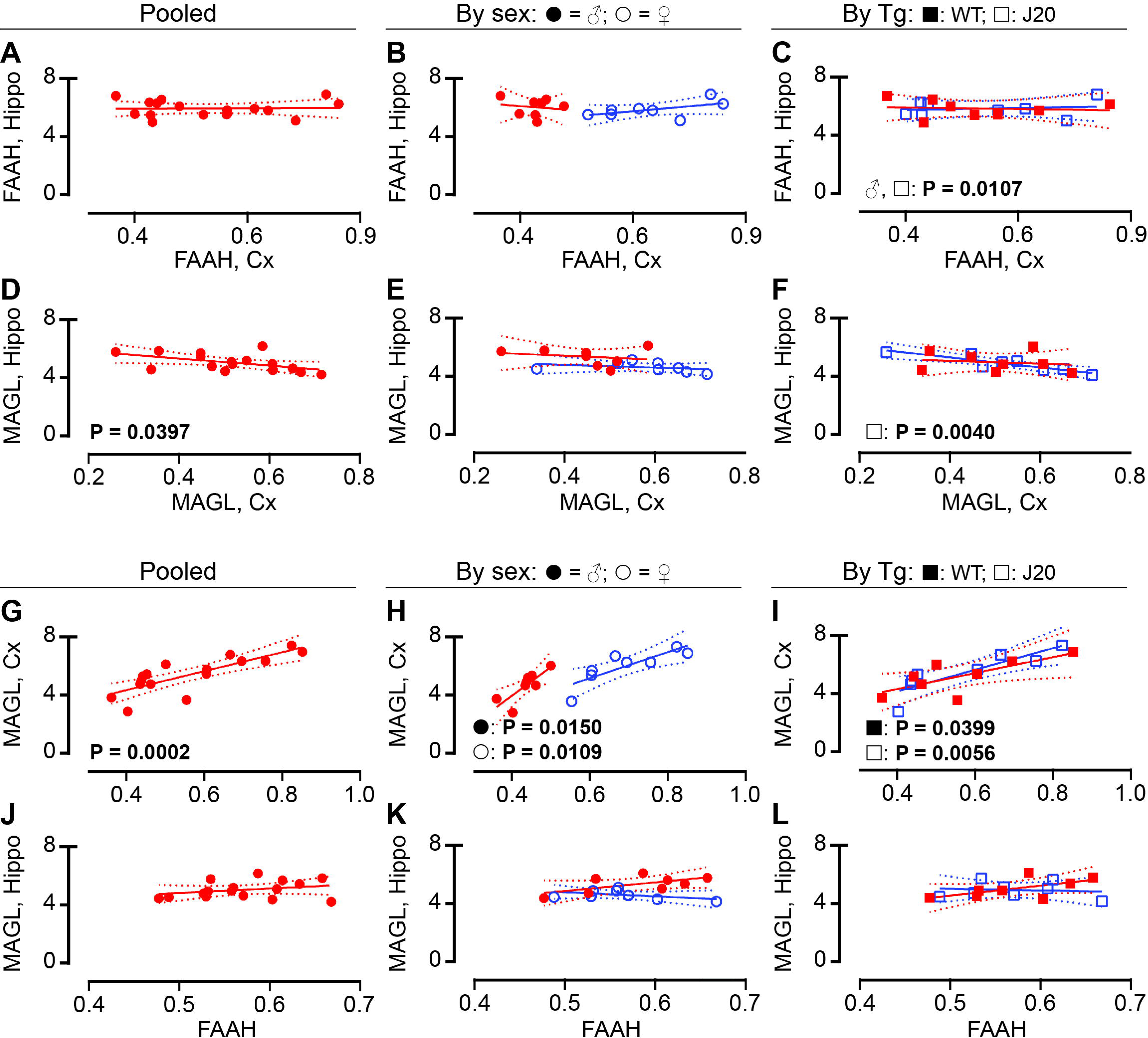
Inter- and intra-region association between FAAH and MAGL in wildtype and J20 mouse brain. Linear regression was used to explore whether any association existed between cortical and hippocampal FAAH expression when data were (**A**) pooled, (**B**) separated by sex alone [▪: male; ○: female], or (**C**) separated by genotype [′: WT; ≤: J20]. (**D-F**) Intra-regional association in MAGL expression was investigated using similar stratification. Within the cortex, associations between FAAH and MAGL level were observed when the data (**G**) were pooled, (**H**) separated by sex, or (**I**) separated by gentopyte. (**J-L**) There were no associations between FAAH and MAGL expression in the corresponding hippocampal asamples. ‘Pooled’ data refers to data being tested regardless of sex or genotype. Only those associations that are significant (*P* ≤ 0.05) are highlighted (next to their grouping symbol).

Within the cortex, there were very strong associations between FAAH and MAGL expression whether the data were pooled [P = 0.0002] (**Figure 7G**) or stratified by sex [male: P = 0.0150; female: P = 0.0109] (**Figure 7H**) or by genotype [WT: P = 0.0399; J20: P = 0.0056] (**Figure 7I**). The association in the WT mouse was driven by female mice [male: P = 0.0834; female: P = 0.0084], whereas any association in the J20 mouse was lost when separated by sex. There were no associations between FAAH and MAGL observed in the corresponding hippocampal samples, regardless of stratification (**Figure 7J-L)**.

#### Alignment of mouse CB1Rs with FAAH or MAGL levels based on stratification by genotype and/or by sex

We did not observe any alignment between CB1R-60 and FAAH or MAGL in mouse cortex (**Figure 8A-C**). Any alignment between CB1Rs and either FAAH or MAGL in mouse brain involved almost exclusively the CB1R-47 and CB1R-37 immunoreactive bands. In cortical samples, FAAH aligns with CB1R-47 in female mice when data is stratified by sex alone mice [male: P = 0.4841; female: P = 0.0016] (**Figure 8B**) and in WT samples when data are stratified by genotype [WT: P = 0.0307; J20: P = 0.7466] (**Figure 8C**). Interestingly, FAAH aligns with CB1R-47 in female J20 samples when data are stratified for sex-by-genotype [male J20: P = 0.5288; female J20: P = 0.0205]. In contrast, FAAH aligns with CB1R-37 exclusively in male mice. Expression of the two proteins is strongly associated if the data are pooled [P < 0.0001] (**Figure 8G**) and in male mice if stratified by sex alone [male: P = 0.0032; female: P = 0.2071] (**Figure 8H**). There are strong associations in both WT and J20 mice [WT: P = 0.0007; J20: P = 0.0086] (**Figure 8I**), which is driven by males in both genotypes [male WT: P = 0.0368; male J20: P = 0.0271 versus female WT: P = 0.1664; female J20: P = 0.5868]. There were no associations between FAAH and CB1Rs in hippocampal samples (**SUPPL Table 26**).

**Figure 8.**
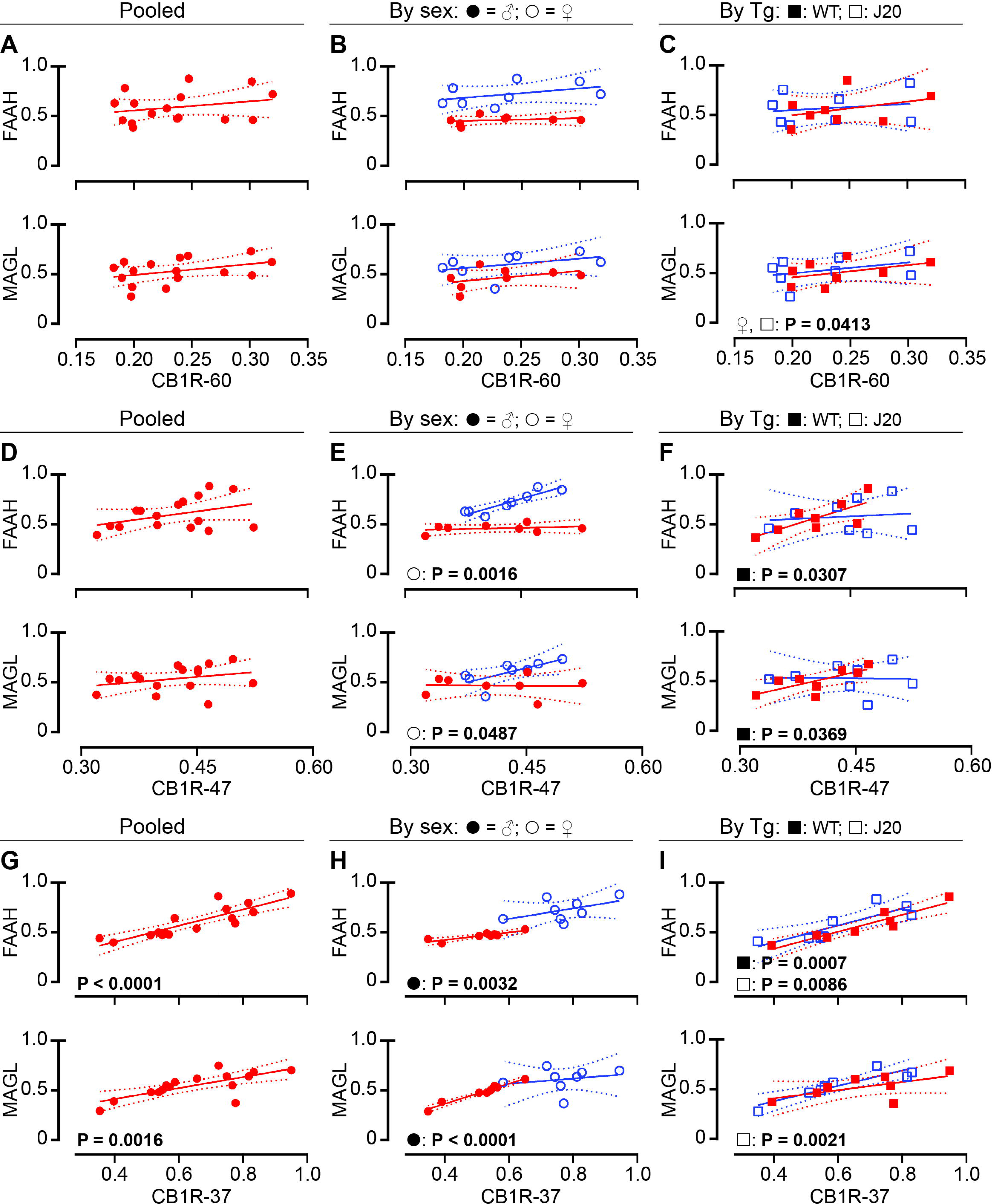
Associations between CB1Rs and either FAAH and MAGL in wildtype and J20 mouse cortex. Linear regression was used to explore whether any association existed between expression of either FAAH (*top row*) or MAGL (*bottom row*) and expression of CB1R-60, CB1R-47, or CB1R-37. Respeticve regession analysis was preformed using (**A, D, G**) pooled data, or (**B, E, H**) data separated by sex alone [▪: male; ○: female], or (**C, F, I**) separated by genotype [′: WT; ≤: J20]. ‘Pooled’ data refers to data being tested regardless of sex or genotype. Only those associations that are significant (*P* ≤ 0.05) are highlighted (next to their grouping symbol).

In cortical samples, MAGL aligns moderately with CB1R-60 in female J20 mice [male: P = 0.5121; female: P = 0.0413] (included in **Figure 8C**, lower panel). MAGL also associates with CB1R-47 when data is stratified by sex alone mice [male: P = 0.9360; female: P = 0.0487] (**Figure 8E**, lower panel) and in WT samples when data are stratified by genotype [WT: P = 0.0369; J20: P = 0.9443] (**Figure 8F**, lower panel). As with the FAAH data, MAGL also aligns with CB1R-37 exclusively in male mice. Expression of the MAGL and CB1R-37 is strongly associated if the data are pooled [P = 0.0016] (**Figure 8G**, lower panel) and in male mice if stratified by sex alone [male: P < 0.0001; female: P = 0.5847] (**Figure 8H**). There was a strong association in between MAGL and CB1R-37 based on genotype [WT: P = 0.1147; J20: P = 0.0021] (**Figure 8I**), with additional analyses indicating males differing from females in both genotypes [male WT: P = 0.0120; male J20: P = 0.0059 versus female WT: P = 0.4583; female J20: P = 0.5292]. Finally, we observed sporadic associations between MAGL and CB1Rs in mouse hippocampal tissues. For example, MAGL associates with CB1R-60 in females [male: P = 0.0846; female: P = 0.0369] and in WT samples [WT: P = 0.0442; J20: P = 0.5100] **(SUPPL Table 27).**

## Discussion

Most cases (>90%) of AD are diagnosed after 65 years of age. Although age and sex are two of the primary risk factors for this late-onset form of AD (LOAD), the cause is largely unknown [43]. Any diagnosis is often inferred from a review of the symptoms (*e.g.,* changes in memory, cognition, behaviour, and functional skills) but can only be confirmed by a post-mortem examination of the brain for histopathological markers. Available medications are not disease-modifying (*e.g.,* they target symptoms but do not reverse neuropathology or the deficits in cognition and memory) and any benefit is diminished within 3-6 months of treatment [76]. Thus, new management strategies are always under review. Components of the ECS, whether it be the CB1R or one of the enzymes, *e.g.* FAAH or MAGL that are involved in regulating levels of the endocannabinoids AEA or 2-AG, respectively, have been implicated in aspects of the disease course and form the basis for numerous reviews on the topic [37, 39, 77–79]. These reviews highlight studies in preclinical models that would support strategies targeting the ECS in AD, with clear benefit to reducing plaque and tau pathology, or improving behavioral, immunological, and cognitive phenotypes. However, these same reviews also highlight that very few, if any, of the strategies benefiting these preclinical models translate effectively to clinical AD. While a meta-analysis found little to no benefit of cannabinoids in AD [80], our own systematic review suggests that cannabinoids may be better suited to provide relief from behavioral and psychological symptoms of dementia, without any overt effect on cognitive phenotypes [81]. In the absence of any information on medication history or cannabinoid usage in the donor casefile, it is impossible to determine how much of the variability we observed in our human samples may be attributed to pharmacotherapeutics. ECS targeting and therapeutic response in AD/dementia is further complicated by influences of sex [82] and *APOE* ε4 status [83, 84] on outcomes.

Study of the ECS relies heavily on relative expression of mRNA transcript levels, yet this may not a reliable practice as *CNR1* mRNA levels may not equate to CB1R expression levels in either control or AD brain [85] or in many other neuropathologies [86]. Our observations confirm the lack of change in *CNR1* mRNA expression in AD samples [85]. Other means of quantifying CB1R expression and function rely on ligand-receptor binding or GTPγS binding, or on immunohistochemical detection, which may not capture all of the information available and often assume a single CB1R entity. Yet this could affect interpretation of data as the literature does refer to multiple anti-CB1R immunodetectable bands, including a 60-kDa band that represents a glycosylated (putatively active) CB1R and a 47-kDa band that migrates at the expected molecular weight of the deduced amino acid sequence for the protein-coding sequence of the *CNR1* gene (*e.g*., without any post-translational modifications) [74]. A ∼37-kDa anti-CB1R signal is proposed to be a putative splice variant [75]. *CNR1* splice variants have been shown to have a higher affinity for 2-AG (*versus* AEA), which appears to act as an inverse agonist on these variants (based on GTPγS activity) [87]. Elsewhere, the expression of *CNR1* variants has been associated with neuropsychiatric diseases such as schizophrenia, bipolar disorder, and major depression [73], all of which present with cognitive phenotypes. The presence of all three bands in our samples provide the means for a broader interpretation of CB1R function during AD. For example, the only change we observe in the mean relative expression levels (compared to normal controls) of any of the CB1R species is the increase in glycosylated/activated CB1R-60 in EOAD and LOAD hippocampal samples. This would corroborate reports that activation of the CB1R in hippocampus alters GABA availability and hinders long-term potentiation, thereby impairing synaptic strength and memory consolidation (reviewed in [88]). The general absence of change in CB1R expression (or in FAAH and MAGL expression) in our AD sample set was intriguing, but not unique as this observation corroborates other reports on the variability in ECS marker expression in AD samples (reviewed in [4, 37, 39, 77–79]). With the exception of an increase in mean cortical CB1R-60 expression in female carriers of the *APOE* ε4 allele (independent of diagnosis), our analyses also did not reveal any overt differences in mean relative expression of markers of the ECS based strictly on the donor’s sex or *APOE* ε4 status (in either region). Interestingly, previous work from our group using these same autopsy samples found clear changes in mean relative expression of markers of other systems, several of which were strongly dependent on sex or *APOE* ε4 status of the donour; these included diverse markers of the serotonin synapse [89], monoamine oxidases A and B [90], and, of course, markers of AD-related amyloid pathology, including the Aβ peptide and related secretases [63, 91].

While there were no overt changes in mean relative expression of markers of the ECS, regression analysis revealed a different scenario. We observed association between the expression of the various CB1R bands, but these were restricted to control samples when the data were stratified by ‘diagnosis’ and far stronger associations were revealed when the data were stratified simply by *APOE* ε4 status or sex alone. In most of these stratifications the association was carried primarily by samples from female donors. Although it is well known that sex [61] and the *APOE* ε4 allele can influence risk of LOAD and amyloid burden (particularly in females) [62, 63], there is a paucity of data regarding the influence of these two major risk factors for AD on regulation of the ECS. For example, there are reports of cannabidiol providing some relief on β-amyloid burden in the male *APP*_Swe_/*PSEN1*ρEx9 mouse [92], but little relief in the male TAU58/2 model of tauopathy and dementia [93], whereas autoradiography reveals a decrease in striatal CB1R density in the female *APP*_Swe_/*PSEN1*(L166P) mouse [94]. A PET study found no difference in CB1R density between AD patients and cognitively intact controls, regardless of *APOE* ε4 status; unfortunately, the test cohort was limited in size, so male and female data were pooled [51]. The few reports of an influence of *APOE* ε4 status on the ECS in AD/dementia centers on cognitive phenotypes in older individuals with metabolic syndrome [84] or dampening of any beneficial effect of exercise on memory in young carriers of the risk allele [83]. Any reports of an influence of *APOE* ε4 status on the ECS enzymes centers on FAAH in stroke [95] and atherosclerosis [96] and MAGL in CB2R-sensitive atherosclerosis [97].

Elsewhere, it was shown that 2-AG concentrations were negatively correlated with baseline cognition and cognitive changes in men and carriers of the *APOE* ε4 allele, while in women, cognitive changes correlated with the abundance of AEA [84]. Interestingly, our observations indicate that levels of CB1Rs (*e.g*., the CB1R-47 or CB1R-37) associate strongly with MAGL expression (and only sporadically with FAAH) and this is particularly evident when the data are stratified by *APOE* ε4 status or sex alone. Again, there appears to be a stronger female prevalence in these observations. The fact that expression of CB1R(47/37) align with MAGL (which hydrolyzes almost 85% of the brain’s 2-AG [98]) suggests that 2-AG plays a significant role in regulating *CNR1* translation and CB1R availability, but not its activation (if we continue to view CB1R-60 as the glycosylated/activated form). The fact that we do not observe any association between CB1Rs and MAGL when the data are stratified by diagnosis suggests *a priori* that the MAGL system may be more sensitive to AD-related cellular phenotypes.

Any positive associations between CB1R species, whether they are post-translational modified (such as the glycosylated CB1R-60) or not, in addition to the observation that expression levels of the different species do not appear to be mutually exclusive (*e.g*. the expression of CB1R-47 is not promoted at the expense of CB1R-37 or *vice versa*), suggests a concerted regulation of both CB1R transcription and function. The fact that these associations are evident almost exclusively in cortical samples and when data are stratified by *APOE* ε4 status or sex alone (but not when stratified for diagnosis), further suggests it is an inherent character of the cortex’s ECS and that its disruption may reflect progression of an AD cortical phenotype.

In clear contrast, the screening of mouse tissues (cortical and corresponding hippocampal samples) revealed intra- and inter-regional alignment between the expression of the different CB1R species as well as between FAAH and/or MAGL. Furthermore, in these tissues, expression levels of CB1Rs (particularly CB1R-47 and CB1R-37) associated with both FAAH and MAGL in both regions and were driven by both sexes, with a potential for slightly more representation in male samples. This does not support the observations made in our human samples, which carried a female bias. It is always a possibility that some of these divergent species-based observations could reflect the multiple subtypes of AD—based on RNASeq [99], CSF proteomics [100], and MRI/imaging [101]—involving distinct signaling pathways and/or pathological markers (*e.g.* amyloid burden, tau phosphorylation/tangles). However, such clinical heterogeneity would not explain why human CB1R expression would align exclusively with MAGL in a region-dependent manner, whereas mouse CB1R expression would align with MAGL as well as FAAH and do so in both the cortex and hippocampus. Furthermore, such heterogeneity would not provide an obvious explanation as to why sex-dependent alignment differs between the two species. As the mouse phenotypes are not affected by the hAPP_Swe/Ind_ transgene, this adds to the discrepancy between species as it suggests a less selective regulation of presynaptic CB1R by AEA or 2-AG in the mouse brain that is not overtly impacted by disease-causing (Aβ-promoting) mutations in APP. The fact that CB1Rs, FAAH, and MAGL appear to be influenced by genotype in, for example, 3xTg-AD [60], 5xFAD [102], or APP_Swe_/PS1ΔE9 [103] mice, all of which carry some familial AD-related variant of the *PSEN1* gene, suggests that mutations in presenilin-1 may exert more influence on ECS phenotypes than any AD-related mutations in APP.

The literature already supports differences between the human and rodent (mouse/rat) ECS. For example, species differences exist with regards to CB1R distribution and its affinity for ligands [30], with greater densities of the CB1R in regions that align with cognition (*e.g*. frontal cortex) and memory (*e.g*. hippocampus) in the human brain, but in regions that align with motor function (*e.g*. cerebellum and caudate putamen) in the rat. Similarly, the human and rodent enzymes involved in regulating endocannabinoid levels, *e.g*. FAAH and MAGL, also differ in distribution and function. While extrapolating the complete range of ECS outcomes (*e.g*., pathological, behavioural, cognitive) from AD-related mouse models to the clinical context may not be feasible, these same models could still provide critical insight on specific phenotypes associated with disease course. For example, the AD-related APP23 mouse strain on a CB1R^−/−^ background carried a lower amyloid plaque burden, but increased cognitive deficits, than mice on a CB1R^+/+^ background [104]. Elsewhere, enhanced Aβ clearance from the brain was observed following administration of 2-AG, CB1R agonists, and a MAGL inhibitor, but not with FAAH inhibitors [105]. We are currently investigating whether the changes in CB1R, FAAH, or MAGL align with either Aβ or Tau pathology in these same human and J20 tissue sets.

Pathway effects of FAAH and MAGL inhibitors suggest roles for FAAH in stress paradigms and roles for MAGL in inflammation in relation to health and disease, with opposing effects of AEA and 2-AG on decision-making and cognitive flexibility (reviewed in [106]). Although MAGL is expressed in proximity to CB1Rs in brain, during the course of AD MAGL (but not FAAH) may not be recruited to its ‘proper’ localization within the cell, thereby affecting 2-AG termination and signaling [50]. In the rodent brain, both FAAH [107] and MAGL [108] are shown to co-distribute with CB1R, whereas in the human brain the expression of the two enzymes appear to differ slightly, allowing FAAH to terminate ECS signaling at the site of synthesis and MAGL terminating ECS signaling at the site of action [16].

## Limitations and conclusions

We acknowledge the focus of this report being CB1R limits the interpretation of the role of the ECS in our samples and that 2-AG and AEA interact with numerous proteins other than the CB1R, *e.g.* [109, 110], any one of which could also be contributing to AD-related phenotypes. A statistical limitation of this study and the interpretation of our data is that our random 60-sample set was not sufficiently powered to undertake relevant three-way stratification, i.e. sex-by-*APOE* ε4-by-diagnosis, although being able to compare between two brain regions from the same donor adds a level of validity to our measured outcomes. As with any comparative study of this nature, there is bias that lies with the inherent variability associated with the heterogeneity of human autopsy sample sets *versus* the homogeneity of inbred mouse strains used in preclinical studies of AD. We acknowledge such inherent variability could impact observations; however, we also feel that while CB1R levels align primarily with MAGL in a sex- and region-dependent manner in human brain and are disrupted when data are stratified by diagnosis, the fact that CB1R levels align with both MAGL and FAAH in mouse cortex and hippocampus regardless of stratification (e.g. sex or genotype/diagnosis) supports valid species-dependent regulatory differences in the ECS. Clearly, additional studies will need to determine whether these observations in human samples are generalizable across other pathologies in which the ECS has been implicated, for example, Parkinson’s disease [111], depression [112], or ALS [113], and in corresponding *in vivo* models. Our study clearly demonstrates sex-dependent differences in the mean relative levels of markers of the ECS in WT mice. Yet, we also observe very clear evidence of an alignment of CB1R, FAAH, and MAGL regardless of genotype (*e.g*., WT or J20). It will be important to determine to what extent our observations made in the J20 mouse apply to other models of AD-related pathology and cognitive deficits used to study the ECS, such as the 3xTg-AD mouse [60] and the 5xFAD mouse [102], or more recently, the APP_Swe_/PS1ΔE9 mouse that exhibits significant region- and sex-dependent differences in *Cnr1*, *Faah*, and *Mgll* mRNA transcript levels that, *a priori*, do not appear to always co-distribute [103].

## Author contributions

RMH, JNN, and DM experimental design. RMH, JNN, PRP, HS, LVBG, and DDM data collection and analysis. All authors were involved in editing the manuscript.

## Acknowledgments

Parts of this work were presented at the 41^st^ Annual Meeting of the Canadian College of Neuropsychopharmacology. This work was funded, in part, by the Saskatchewan Research Chair in Alzheimer Disease and Related Dementia, which is funded jointly by the Alzheimer Society of Saskatchewan and the Saskatchewan Health Research Foundation. There are no competing interests.

**Supplementary Figures 1-4:** These figures include information, primarily mean expression levels, in which only sporadic associations (*P* < 0.05) were observed.

**Supplementary Tables 1-27:** These tables include information, primarily results of regression analyses, in which only sporadic associations (*P* < 0.05) were observed.

