## Supplementary material for "Differences in the co-distribution of the Cannabinoid Receptor-1, FAAH, and MAGL in the human and mouse brain could be limiting clinical translation of endocannabinoid outcomes in mouse models of Alzheimer disease": SUPPL Tables 1-27

| Cortex | 60 *vs* 47 | CTL | | EOAD | | LOAD | |
|---|---|---|---|---|---|---|---|
| | | P = 0.063 | | P = 0.182 | | P = 0.581 | |
| | | $R^2$ = 0.142 | | $R^2$ = 0.123 | | $R^2$ = 0.020 | |
| | Diagnosis by sex | Male | Female | Male | Female | Male | Female |
| | | P = 0.537 | P = 0.099 | P = 0.718 | P = 0.706 | P = 0.475 | P = 0.945 |
| | | $R^2$ = 0.039 | $R^2$ = 0.228 | $R^2$ = 0.036 | $R^2$ = 0.022 | $R^2$ = 0.088 | $R^2$ = 0.000 |
| Cortex | 60 *vs* 37 | CTL | | EOAD | | LOAD | |
| | | **P = 0.032** | | P = 0.399 | | P = 0.479 | |
| | | $R^2$ = 0.185 | | $R^2$ = 0.051 | | $R^2$ = 0.032 | |
| | Diagnosis by sex | Male | Female | Male | Female | Male | Female |
| | | P = 0.297 | **P = 0.008** | P = 0.691 | P = 0.508 | P = 0.911 | P = 0.576 |
| | | $R^2$ = 0.108 | $R^2$ = 0.486 | $R^2$ = 0.034 | $R^2$ = 0.065 | $R^2$ = 0.002 | $R^2$ = 0.041 |
| Cortex | 47 *vs* 37 | CTL | | EOAD | | LOAD | |
| | | **P = 0.034** | | **P = 0.011** | | P = 0.169 | |
| | | $R^2$ = 0.180 | | $R^2$ = 0.377 | | $R^2$ = 0.115 | |
| | Diagnosis by sex | Male | Female | Male | Female | Male | Female |
| | | P = 0.379 | **P = 0.000** | P = 0.237 | P = 0.062 | P = 0.078 | P = 0.452 |
| | | $R^2$ = 0.078 | $R^2$ = 0.761 | $R^2$ = 0.265 | $R^2$ = 0.413 | $R^2$ = 0.430 | $R^2$ = 0.073 |

**SUPPL Table 1: Linear regression (by diagnosis): CB1R species *vs* CB1R species within cortex. Significant (P ≤ 0.05) observations, if any, are identified in bold text.**

| | | | | | | | |
|---|---|---|---|---|---|---|---|
| Hippocampus | 60 *vs* 47 | CTL | | EOAD | | LOAD | |
| | | P = 0.921 | | P = 0.322 | | P = 0.214 | |
| | | $R^2$ = 0.000 | | $R^2$ = 0.075 | | $R^2$ = 0.090 | |
| | Diagnosis by sex | Male | Female | Male | Female | Male | Female |
| | | P = 0.946 | P = 0.999 | P = 0.822 | P = 0.551 | **P = 0.044** | P = 0.638 |
| | | $R^2$ = 0.001 | $R^2$ = 0.000 | $R^2$ = 0.011 | $R^2$ = 0.063 | $R^2$ = 0.591 | $R^2$ = 0.029 |
| Hippocampus | 60 *vs* 37 | CTL | | EOAD | | LOAD | |
| | | P = 0.965 | | P = 0.419 | | P = 0.447 | |
| | | $R^2$ = 0.000 | | $R^2$ = 0.051 | | $R^2$ = 0.039 | |
| | Diagnosis by sex | Male | Female | Male | Female | Male | Female |
| | | P = 0.657 | P = 0.616 | P = 0.523 | P = 0.626 | P = 0.601 | P = 0.952 |
| | | $R^2$ = 0.054 | $R^2$ = 0.027 | $R^2$ = 0.087 | $R^2$ = 0.042 | $R^2$ = 0.059 | $R^2$ = 0.001 |
| Hippocampus | 47 *vs* 37 | CTL | | EOAD | | LOAD | |
| | | P = 0.069 | | P = 0.813 | | P = 0.782 | |
| | | $R^2$ = 0.193 | | $R^2$ = 0.005 | | $R^2$ = 0.079 | |
| | Diagnosis by sex | Male | Female | Male | Female | Male | Female |
| | | P = 0.071 | P = 0.424 | P = 0.089 | P = 0.974 | P = 0.753 | P = 0.786 |
| | | $R^2$ = 0.600 | $R^2$ = 0.065 | $R^2$ = 0.471 | $R^2$ = 0.000 | $R^2$ = 0.022 | $R^2$ = 0.010 |

**SUPPL Table 2: Linear regression (by diagnosis): CB1R species *vs* CB1R species within hippocampus. Significant ($P \leq 0.05$) observations, if any, are identified in bold text.**

| | | | | | | | |
|---|---|---|---|---|---|---|---|
| Cortex *vs* Hippocampus | 60 *vs* 60 | CTL | | EOAD | | LOAD | |
| | | P = 0.745 | | **P = 0.048** | | P = 0.232 | |
| | | $R^2$ = 0.007 | | $R^2$ = 0.288 | | $R^2$ = 0.094 | |
| | Diagnosis by sex | Male | Female | Male | Female | Male | Female |
| | | P = 0.872 | P = 0.855 | P = 0.090 | P = 0.321 | P = 0.886 | P = 0.106 |
| | | $R^2$ = 0.007 | $R^2$ = 0.004 | $R^2$ = 0.469 | $R^2$ = 0.195 | $R^2$ = 0.005 | $R^2$ = 0.293 |
| Cortex *vs* Hippocampus | 47 *vs* 47 | CTL | | EOAD | | LOAD | |
| | | P = 0.427 | | P = 0.327 | | P = 0.271 | |
| | | $R^2$ = 0.043 | | $R^2$ = 0.074 | | $R^2$ = 0.080 | |
| | Diagnosis by sex | Male | Female | Male | Female | Male | Female |
| | | P = 0.802 | P = 0.496 | P = 0.560 | P = 0.368 | P = 0.370 | P = 0.337 |
| | | $R^2$ = 0.018 | $R^2$ = 0.053 | $R^2$ = 0.072 | $R^2$ = 0.137 | $R^2$ = 0.162 | $R^2$ = 0.116 |
| Cortex *vs* Hippocampus | 37 *vs* 37 | CTL | | EOAD | | LOAD | |
| | | P = 0.073 | | **P = 0.003** | | P = 0.642 | |
| | | $R^2$ = 0.199 | | $R^2$ = 0.502 | | $R^2$ = 0.015 | |
| | Diagnosis by sex | Male | Female | Male | Female | Male | Female |
| | | P = 0.079 | P = 0.581 | P = 0.564 | **P = 0.003** | P = 0.865 | P = 0.418 |
| | | $R^2$ = 0.579 | $R^2$ = 0.035 | $R^2$ = 0.071 | $R^2$ = 0.786 | $R^2$ = 0.006 | $R^2$ = 0.083 |

**SUPPL Table 3: Linear regression (by diagnosis): CB1R species *between* cortex and hippocampus. Significant ($P \leq 0.05$) observations, if any, are identified in bold text.**

| | | | | | |
|---|---|---|---|---|---|
| Hippocampus | 60 *vs* 47 | Male (-ε4) | Male (+ε4) | Female (-ε4) | Female (+ε4) |
| | | P = 0.286 | P = 0.098 | P = 0.865 | P = 0.681 |
| | | $R^2$ = 0.222 | $R^2$ = 0.229 | $R^2$ = 0.003 | $R^2$ = 0.013 |
| Hippocampus | 60 *vs* 37 | Male (-ε4) | Male (+ε4) | Female (-ε4) | Female (+ε4) |
| | | **P = 0.018** | P = 0.953 | P = 0.847 | P = 0.575 |
| | | $R^2$ = 0.706 | $R^2$ = 0.000 | $R^2$ = 0.002 | $R^2$ = 0.023 |
| Hippocampus | 47 *vs* 37 | Male (-ε4) | Male (+ε4) | Female (-ε4) | Female (+ε4) |
| | | P = 0.437 | P = 0.073 | P = 0.313 | P = 0.371 |
| | | $R^2$ = 0.125 | $R^2$ = 0.264 | $R^2$ = 0.085 | $R^2$ = 0.057 |

**SUPPL Table 4: Association between CB1R species within hippocampus (by *APOE* ε4-by-sex). Significant (P ≤ 0.05) observations, if any, are identified in bold text.**

| | | | |
|---|---|---|---|
| Hippocampus | 60 *vs* 47 | Male | Female |
| | | P = 0.117 | P = 0.164 |
| | | $R^2$ = 0.131 | $R^2$ = 0.068 |
| Hippocampus | 60 *vs* 37 | Male | Female |
| | | P = 0.261 | P = 0.305 |
| | | $R^2$ = 0.070 | $R^2$ = 0.038 |
| Hippocampus | 47 *vs* 37 | Male | Female |
| | | P = 0.102 | P = 0.675 |
| | | $R^2$ = 0.141 | $R^2$ = 0.006 |

**SUPPL Table 5: Association between CB1R species within hippocampus (by sex alone). Significant ($P \leq 0.05$) observations, if any, are identified in bold text.**

| Cortex *vs* Hippocampus | 60 *vs* 60 | By sex | | *APOE* -ε4 | | *APOE* +ε4 | |
|---|---|---|---|---|---|---|---|
| | | Male | Female | Male | Female | Male | Female |
| | | P = 0.129<br>$R^2$ = 0.123 | P = 0.086<br>$R^2$ = 0.109 | P = 0.605<br>$R^2$ = 0.057 | P = 0.074<br>$R^2$ = 0.262 | P = 0.129<br>$R^2$ = 0.197 | P = 0.472<br>$R^2$ = 0.041 |
| **Cortex *vs* Hippocampus** | **47 *vs* 47** | **By sex** | | ***APOE* -ε4** | | ***APOE* +ε4** | |
| | | Male | Female | Male | Female | Male | Female |
| | | P = 0.428<br>$R^2$ = 0.035 | P = 0.849<br>$R^2$ = 0.001 | P = 0.366<br>$R^2$ = 0.165 | P = 0.967<br>$R^2$ = 0.000 | P = 0.560<br>$R^2$ = 0.032 | P = 0.933<br>$R^2$ = 0.000 |
| **Cortex *vs* Hippocampus** | **37 *vs* 37** | **By sex** | | ***APOE* -ε4** | | ***APOE* +ε4** | |
| | | Male | Female | Male | Female | Male | Female |
| | | P = 0.367<br>$R^2$ = 0.045 | **P = 0.025**<br>$R^2$ = 0.173 | P = 0.634<br>$R^2$ = 0.049 | P = 0.742<br>$R^2$ = 0.010 | P = 0.576<br>$R^2$ = 0.029 | **P = 0.008**<br>$R^2$ = 0.409 |

**SUPPL Table 6: Linear regression (by sex or by *APOE* ε4 status): CB1R species between cortex and hippocampus. Significant (P ≤ 0.05) observations, if any, are identified in bold text.**

| Cortex *vs* Hippocampus | 60 *vs* 60 | By sex | | *APOE* -ε4 | | *APOE* +ε4 | |
|---|---|---|---|---|---|---|---|
| | | Male | Female | Male | Female | Male | Female |
| | | P = 0.129<br>$R^2$ = 0.123 | P = 0.086<br>$R^2$ = 0.109 | P = 0.605<br>$R^2$ = 0.057 | P = 0.074<br>$R^2$ = 0.262 | P = 0.129<br>$R^2$ = 0.197 | P = 0.472<br>$R^2$ = 0.041 |
| Cortex *vs* Hippocampus | 47 *vs* 47 | By sex | | *APOE* -ε4 | | *APOE* +ε4 | |
| | | Male | Female | Male | Female | Male | Female |
| | | P = 0.428<br>$R^2$ = 0.035 | P = 0.849<br>$R^2$ = 0.001 | P = 0.366<br>$R^2$ = 0.165 | P = 0.967<br>$R^2$ = 0.000 | P = 0.560<br>$R^2$ = 0.032 | P = 0.933<br>$R^2$ = 0.000 |
| Cortex *vs* Hippocampus | 37 *vs* 37 | By sex | | *APOE* -ε4 | | *APOE* +ε4 | |
| | | Male | Female | Male | Female | Male | Female |
| | | P = 0.367<br>$R^2$ = 0.045 | **P = 0.025**<br>$R^2$ = 0.173 | P = 0.634<br>$R^2$ = 0.049 | P = 0.742<br>$R^2$ = 0.010 | P = 0.576<br>$R^2$ = 0.029 | **P = 0.008**<br>$R^2$ = 0.409 |

**SUPPL Table 7: Linear regression (by sex or by *APOE* ε4 status): CB1R species between cortex and hippocampus. Significant (P ≤ 0.05) observations, if any, are identified in bold text.**

| | | | | | |
|---|---|---|---|---|---|
| Hippocampus | 60 *vs* 47 | Male (-ε4) | Male (+ε4) | Female (-ε4) | Female (+ε4) |
| | | $P = 0.286$<br>$R^2 = 0.222$ | $P = 0.098$<br>$R^2 = 0.229$ | $P = 0.865$<br>$R^2 = 0.003$ | $P = 0.681$<br>$R^2 = 0.013$ |
| Hippocampus | 60 *vs* 37 | Male (-ε4) | Male (+ε4) | Female (-ε4) | Female (+ε4) |
| | | **P = 0.018**<br>$R^2 = 0.706$ | $P = 0.953$<br>$R^2 = 0.000$ | $P = 0.847$<br>$R^2 = 0.002$ | $P = 0.575$<br>$R^2 = 0.023$ |
| Hippocampus | 47 *vs* 37 | Male (-ε4) | Male (+ε4) | Female (-ε4) | Female (+ε4) |
| | | $P = 0.437$<br>$R^2 = 0.125$ | $P = 0.073$<br>$R^2 = 0.264$ | $P = 0.313$<br>$R^2 = 0.085$ | $P = 0.371$<br>$R^2 = 0.057$ |

**SUPPL Table 8: Association between CB1R species within hippocampus (by APOE ε4-by-sex). Significant ($P \leq 0.05$) observations, if any, are identified in bold text.**

| Cortex | FAAH vs MAGL | CTL | | EOAD | | LOAD | |
|---|---|---|---|---|---|---|---|
| | | P = 0.132 | | P = 0.704 | | P = 0.575 | |
| | | $R^2$ = 0.096 | | $R^2$ = 0.011 | | $R^2$ = 0.020 | |
| | by sex | Male | Female | Male | Female | Male | Female |
| | | P = 0.335 | P = 0.246 | P = 0.075 | P = 0.416 | P = 0.862 | P = 0.476 |
| | | $R^2$ = 0.093 | $R^2$ = 0.120 | $R^2$ = 0.501 | $R^2$ = 0.097 | $R^2$ = 0.005 | $R^2$ = 0.066 |

| Hippocampus | FAAH vs MAGL | CTL | | EOAD | | LOAD | |
|---|---|---|---|---|---|---|---|
| | | P = 0.119 | | P = 0.079 | | P = 0.273 | |
| | | $R^2$ = 0.145 | | $R^2$ = 0.218 | | $R^2$ = 0.085 | |
| | by sex | Male | Female | Male | Female | Male | Female |
| | | **P = 0.035** | P = 0.260 | P = 0.077 | P = 0.280 | P = 0.678 | P = 0.276 |
| | | $R^2$ = 0.710 | $R^2$ = 0.125 | $R^2$ = 0.496 | $R^2$ = 0.190 | $R^2$ = 0.048 | $R^2$ = 0.146 |

**SUPPL Table 9: Linear regression (by diagnosis): MAGL *vs* FAAH within cortex or hippocampus. Significant (P ≤ 0.05) observations, if any, are identified in bold text.**

| Cortex *vs* Hippocampus | FAAH *vs* FAAH | CTL | | EOAD | | LOAD | |
|---|---|---|---|---|---|---|---|
| | | **P = 0.029** | | P = 0.824 | | P = 0.343 | |
| | | $R^2$ = 0.280 | | $R^2$ = 0.005 | | $R^2$ = 0.060 | |
| | by sex | Male | Female | Male | Female | Male | Female |
| | | P = 0.316 | P = 0.111 | P = 0.449 | P = 0.915 | P = 0.834 | P = 0.408 |
| | | $R^2$ = 0.247 | $R^2$ = 0.258 | $R^2$ = 0.119 | $R^2$ = 0.002 | $R^2$ = 0.010 | $R^2$ = 0.087 |
| Cortex *vs* Hippocampus | MAGL vs MAGL | CTL | | EOAD | | LOAD | |
| | | P = 0.674 | | P = 0.192 | | P = 0.599 | |
| | | $R^2$ = 0.012 | | $R^2$ = 0.127 | | $R^2$ = 0.020 | |
| | by sex | Male | Female | Male | Female | Male | Female |
| | | P = 0.519 | P = 0.644 | P = 0.161 | P = 0.915 | P = 0.505 | P = 0.474 |
| | | $R^2$ = 0.111 | $R^2$ = 0.025 | $R^2$ = 0.351 | $R^2$ = 0.002 | $R^2$ = 0.118 | $R^2$ = 0.066 |

**SUPPL Table 10: Linear regression (by diagnosis): MAGL *vs* FAAH between cortex and hippocampus. Significant (P ≤ 0.05) observations, if any, are identified in bold text.**

| Cortex *vs* Hippocampus | FAAH | *APOE* -ε4 | *APOE* +ε4 | *APOE* -ε4 by sex | | *APOE* +ε4 by sex | |
|---|---|---|---|---|---|---|---|
| | | | | Male | Female | Male | Female |
| | | **P = 0.011** | P = 0.949 | P = 0.089 | P = 0.087 | P = 0.475 | P = 0.464 |
| | | $R^2$ = 0.310 | $R^2$ = 0.000 | $R^2$ = 0.470 | $R^2$ = 0.243 | $R^2$ = 0.066 | $R^2$ = 0.039 |
| Cortex *vs* Hippocampus | MAGL | *APOE* -ε4 | *APOE* +ε4 | *APOE* -ε4 by sex | | *APOE* +ε4 by sex | |
| | | | | Male | Female | Male | Female |
| | | P = 0.297 | P = 0.431 | P = 0.689 | P = 0.666 | P = 0.689 | P = 0.500 |
| | | $R^2$ = 0.060 | $R^2$ = 0.024 | $R^2$ = 0.034 | $R^2$ = 0.018 | $R^2$ = 0.017 | $R^2$ = 0.033 |

**SUPPL Table 11: Linear regression (by *APOE* ε4 status): FAAH or MAGL between cortex and hippocampus. Significant (P ≤ 0.05) observations, if any, are identified in bold text.**

| Cortex *vs* Hippocampus | FAAH | Male | Female |
|---|---|---|---|
| | | P = 0.077 | P = 0.285 |
| | | $R^2$ = 0.194 | $R^2$ = 0.042 |

| Cortex *vs* Hippocampus | MAGL | Male | Female |
|---|---|---|---|
| | | P = 0.356 | P = 0.380 |
| | | $R^2$ = 0.050 | $R^2$ = 0.029 |

**SUPPL Table 12: Linear regression (by sex alone): FAAH or MAGL between cortex and hippocampus. Significant ($P \leq 0.05$) observations, if any, are identified in bold text.**

| Cortex | CB1R-60 *vs* FAAH | CTL | | EOAD | | LOAD | |
|---|---|---|---|---|---|---|---|
| | | P = 0.271 | | P = 0.225 | | P = 0.827 | |
| | | $R^2$ = 0.052 | | $R^2$ = 0.103 | | $R^2$ = 0.003 | |
| | by sex | Male | Female | Male | Female | Male | Female |
| | | P = 0.849 | P = 0.167 | P = 0.205 | P = 0.223 | P = 0.455 | P = 0.875 |
| | | $R^2$ = 0.004 | $R^2$ = 0.166 | $R^2$ = 0.298 | $R^2$ = 0.204 | $R^2$ = 0.096 | $R^2$ = 0.003 |
| Cortex | CB1R-47 *vs* FAAH | CTL | | EOAD | | LOAD | |
| | | P = 0.707 | | P = 0.088 | | P = 0.097 | |
| | | $R^2$ = 0.006 | | $R^2$ = 0.194 | | $R^2$ = 0.163 | |
| | by sex | Male | Female | Male | Female | Male | Female |
| | | P = 0.652 | P = 0.817 | P = 0.215 | P = 0.176 | P = 0.920 | **P = 0.011** |
| | | $R^2$ = 0.021 | $R^2$ = 0.005 | $R^2$ = 0.287 | $R^2$ = 0.245 | $R^2$ = 0.002 | $R^2$ = 0.575 |
| Cortex | CB1R-37 *vs* FAAH | CTL | | EOAD | | LOAD | |
| | | P = 0.347 | | P = 0.746 | | P = 0.950 | |
| | | $R^2$ = 0.039 | | $R^2$ = 0.008 | | $R^2$ = 0.000 | |
| | by sex | Male | Female | Male | Female | Male | Female |
| | | P = 0.261 | P = 0.774 | P = 0.783 | P = 0.832 | P = 0.767 | P = 0.848 |
| | | $R^2$ = 0.124 | $R^2$ = 0.008 | $R^2$ = 0.017 | $R^2$ = 0.007 | $R^2$ = 0.016 | $R^2$ = 0.005 |

**SUPPL Table 13: Linear regression (by diagnosis): Cortex FAAH *vs* CB1R species. Significant (P ≤ 0.05) observations, if any, are identified in bold text.**

| Hippocampus | CB1R-60 *vs* FAAH | CTL | | EOAD | | LOAD | |
|---|---|---|---|---|---|---|---|
| | | P = 0.650 | | P = 0.766 | | P = 0.908 | |
| | | $R^2$ = 0.013 | | $R^2$ = 0.008 | | $R^2$ = 0.000 | |
| | by sex | Male | Female | Male | Female | Male | Female |
| | | P = 0.344 | P = 0.965 | P = 0.664 | P = 0.199 | P = 0.765 | P = 0.411 |
| | | $R^2$ = 0.223 | $R^2$ = 0.000 | $R^2$ = 0.041 | $R^2$ = 0.305 | $R^2$ = 0.020 | $R^2$ = 0.086 |
| Hippocampus | CB1R-47 *vs* FAAH | CTL | | EOAD | | LOAD | |
| | | **P = 0.022** | | P = 0.381 | | P = 0.762 | |
| | | $R^2$ = 0.287 | | $R^2$ = 0.060 | | $R^2$ = 0.006 | |
| | by sex | Male | Female | Male | Female | Male | Female |
| | | P = 0.116 | P = 0.059 | P = 0.945 | P = 0.152 | P = 0.512 | P = 0.316 |
| | | $R^2$ = 0.499 | $R^2$ = 0.312 | $R^2$ = 0.001 | $R^2$ = 0.310 | $R^2$ = 0.090 | $R^2$ = 0.125 |
| Hippocampus | CB1R-37 *vs* FAAH | CTL | | EOAD | | LOAD | |
| | | P = 0.148 | | P = 0.933 | | P = 0.382 | |
| | | $R^2$ = 0.126 | | $R^2$ = 0.000 | | $R^2$ = 0.051 | |
| | by sex | Male | Female | Male | Female | Male | Female |
| | | P = 0.284 | P = 0.296 | P = 0.657 | P = 0.751 | P = 0.481 | P = 0.574 |
| | | $R^2$ = 0.276 | $R^2$ = 0.109 | $R^2$ = 0.043 | $R^2$ = 0.018 | $R^2$ = 0.104 | $R^2$ = 0.041 |

**SUPPL Table 14: Linear regression (by diagnosis): Hippocampus FAAH *vs* CB1R species. Significant (P ≤ 0.05) observations, if any, are identified in bold text.**

| Cortex | CB1R-60 *vs* MAGL | CTL | | EOAD | | LOAD | |
|---|---|---|---|---|---|---|---|
| | | P = 0.276 | | P = 0.351 | | P = 0.353 | |
| | | $R^2$ = 0.051 | | $R^2$ = 0.062 | | $R^2$ = 0.054 | |
| | by sex | Male | Female | Male | Female | Male | Female |
| | | P = 0.400 | P = 0.507 | P = 0.193 | P = 0.932 | P = 0.294 | P = 0.593 |
| | | $R^2$ = 0.072 | $R^2$ = 0.041 | $R^2$ = 0.311 | $R^2$ = 0.001 | $R^2$ = 0.181 | $R^2$ = 0.037 |
| Cortex | CB1R-47 *vs* MAGL | CTL | | EOAD | | LOAD | |
| | | **P = 0.006** | | P = 0.159 | | P = 0.134 | |
| | | $R^2$ = 0.279 | | $R^2$ = 0.137 | | $R^2$ = 0.135 | |
| | by sex | Male | Female | Male | Female | Male | Female |
| | | **P = 0.002** | P = 0.127 | P = 0.403 | **P = 0.015** | P = 0.392 | P = 0.281 |
| | | $R^2$ = 0.646 | $R^2$ = 0.190 | $R^2$ = 0.143 | $R^2$ = 0.597 | $R^2$ = 0.124 | $R^2$ = 0.143 |
| Cortex | CB1R-37 *vs* MAGL | CTL | | EOAD | | LOAD | |
| | | P = 0.803 | | P = 0.062 | | **P = 0.009** | |
| | | $R^2$ = 0.003 | | $R^2$ = 0.228 | | $R^2$ = 0.359 | |
| | by sex | Male | Female | Male | Female | Male | Female |
| | | P = 0.768 | P = 0.728 | P = 0.495 | P = 0.077 | P = 0.556 | **P = 0.037** |
| | | $R^2$ = 0.009 | $R^2$ = 0.011 | $R^2$ = 0.098 | $R^2$ = 0.380 | $R^2$ = 0.061 | $R^2$ = 0.437 |

**SUPPL Tale 15: Linear regression (by diagnosis): Cortex MAGL *vs* CB1R species. Significant (P ≤ 0.05) observations, if any, are identified in bold text.**

| | | | | | | | |
|---|---|---|---|---|---|---|---|
| Hippocampus | CB1R-60 *vs* MAGL | CTL | | EOAD | | LOAD | |
| | | P = 0.992 | | P = 0.242 | | P = 0.828 | |
| | | $R^2$ = 0.000 | | $R^2$ = 0.112 | | $R^2$ = 0.004 | |
| | by sex | Male | Female | Male | Female | Male | Female |
| | | P = 0.985 | P = 0.647 | P = 0.938 | **P = 0.034** | P = 0.488 | P = 0.529 |
| | | $R^2$ = 0.000 | $R^2$ = 0.022 | $R^2$ = 0.001 | $R^2$ = 0.625 | $R^2$ = 0.127 | $R^2$ = 0.051 |
| Hippocampus | CB1R-47 *vs* MAGL | CTL | | EOAD | | LOAD | |
| | | **P = 0.000** | | P = 0.149 | | P = 0.566 | |
| | | $R^2$ = 0.583 | | $R^2$ = 0.153 | | $R^2$ = 0.024 | |
| | by sex | Male | Female | Male | Female | Male | Female |
| | | **P = 0.005** | **P = 0.027** | P = 0.544 | **P = 0.032** | **P = 0.025** | P = 0.937 |
| | | $R^2$ = 0.883 | $R^2$ = 0.420 | $R^2$ = 0.078 | $R^2$ = 0.561 | $R^2$ = 0.752 | $R^2$ = 0.000 |
| Hippocampus | CB1R-37 *vs* MAGL | CTL | | EOAD | | LOAD | |
| | | P = 0.114 | | P = 0.423 | | P = 0.587 | |
| | | $R^2$ = 0.149 | | $R^2$ = 0.050 | | $R^2$ = 0.022 | |
| | by sex | Male | Female | Male | Female | Male | Female |
| | | P = 0.188 | P = 0.377 | P = 0.346 | P = 0.917 | P = 0.801 | P = 0.681 |
| | | $R^2$ = 0.386 | $R^2$ = 0.079 | $R^2$ = 0.177 | $R^2$ = 0.002 | $R^2$ = 0.018 | $R^2$ = 0.022 |

**SUPPL Table 16: Linear regression (by diagnosis): Hippocampus MAGL *vs* CB1R species. Significant ($P \leq 0.05$) observations, if any, are identified in bold text.**

| Cortex | FAAH *vs* CB1R-60 | *APOE* -ε4 | *APOE* +ε4 | *APOE* -ε4 by sex | | *APOE* +ε4 by sex | |
|---|---|---|---|---|---|---|---|
| | | | | Male | Female | Male | Female |
| | | P = 0.811<br>$R^2$ = 0.002 | P = 0.631<br>$R^2$ = 0.008 | P = 0.771<br>$R^2$ = 0.009 | P = 0.962<br>$R^2$ = 0.000 | P = 0.210<br>$R^2$ = 0.118 | P = 0.545<br>$R^2$ = 0.027 |
| Cortex | FAAH *vs* CB1R-47 | *APOE* -ε4 | *APOE* +ε4 | *APOE* -ε4 by sex | | *APOE* +ε4 by sex | |
| | | | | Male | Female | Male | Female |
| | | P = 0.168<br>$R^2$ = 0.072 | P = 0.168<br>$R^2$ = 0.065 | P = 0.174<br>$R^2$ = 0.177 | P = 0.296<br>$R^2$ = 0.078 | P = 0.567<br>$R^2$ = 0.026 | P = 0.108<br>$R^2$ = 0.174 |
| Cortex | FAAH *vs* CB1R-37 | *APOE* -ε4 | *APOE* +ε4 | *APOE* -ε4 by sex | | *APOE* +ε4 by sex | |
| | | | | Male | Female | Male | Female |
| | | P = 0.880<br>$R^2$ = 0.001 | P = 0.609<br>$R^2$ = 0.009 | P = 0.571<br>$R^2$ = 0.033 | P = 0.572<br>$R^2$ = 0.023 | P = 0.153<br>$R^2$ = 0.151 | P = 0.761<br>$R^2$ = 0.007 |

**SUPPL Table 17: Linear regression (by *APOE* ε4 status): FAAH vs CB1R species within cortex. Significant (P ≤ 0.05) observations, if any, are identified in bold text.**

| | | | |
|---|---|---|---|
| Cortex | FAAH *vs* CB1R-60 | Male | Female |
| | | P = 0.276<br>$R^2$ = 0.047 | P = 0.879<br>$R^2$ = 0.000 |
| Cortex | FAAH *vs* CB1R-47 | Male | Female |
| | | P = 0.182<br>$R^2$ = 0.070 | **P = 0.049**<br>$R^2$ = 0.122 |
| Cortex | FAAH *vs* CB1R-37 | Male | Female |
| | | P = 0.185<br>$R^2$ = 0.069 | P = 0.588<br>$R^2$ = 0.010 |

**SUPPL Table 18: Association between CB1R species within hippocampus (by sex alone). Significant ($P \leq 0.05$) observations, if any, are identified in bold text.**

| Hippocampus | FAAH *vs* CB1R-60 | *APOE* -ε4 | *APOE* +ε4 | *APOE* -ε4 by sex | | *APOE* +ε4 by sex | |
|---|---|---|---|---|---|---|---|
| | | | | Male | Female | Male | Female |
| | | P = 0.123 | P = 0.810 | P = 0.177 | P = 0.281 | P = 0.842 | P = 0.979 |
| | | $R^2$ = 0.121 | $R^2$ = 0.002 | $R^2$ = 0.330 | $R^2$ = 0.096 | $R^2$ = 0.005 | $R^2$ = 0.000 |
| Hippocampus | FAAH *vs* CB1R-47 | *APOE* -ε4 | *APOE* +ε4 | *APOE* -ε4 by sex | | *APOE* +ε4 by sex | |
| | | | | Male | Female | Male | Female |
| | | P = 0.893 | P = 0.524 | P = 0.494 | P = 0.843 | P = 0.3787 | P = 0.929 |
| | | $R^2$ = 0.001 | $R^2$ = 0.015 | $R^2$ = 0.098 | $R^2$ = 0.003 | $R^2$ = 0.098 | $R^2$ = 0.001 |
| Hippocampus | FAAH *vs* CB1R-37 | *APOE* -ε4 | *APOE* +ε4 | *APOE* -ε4 by sex | | *APOE* +ε4 by sex | |
| | | | | Male | Female | Male | Female |
| | | P = 0.117 | P = 0.728 | P = 0.327 | P = 0.228 | P = 0.950 | P = 0.361 |
| | | $R^2$ = 0.124 | $R^2$ = 0.005 | $R^2$ = 0.191 | $R^2$ = 0.119 | $R^2$ = 0.001 | $R^2$ = 0.060 |

**SUPPL Table 19: Linear regression (by *APOE* ε4 status): FAAH vs CB1R species within hippocampus. Significant ($P \leq 0.05$) observations, if any, are identified in bold text.**

| Hippocampus | FAAH *vs* CB1R-60 | *APOE* -ε4 | *APOE* +ε4 | *APOE* -ε4 by sex | | *APOE* +ε4 by sex | |
|---|---|---|---|---|---|---|---|
| | | | | Male | Female | Male | Female |
| | | P = 0.123 | P = 0.810 | P = 0.177 | P = 0.281 | P = 0.842 | P = 0.979 |
| | | $R^2$ = 0.121 | $R^2$ = 0.002 | $R^2$ = 0.330 | $R^2$ = 0.096 | $R^2$ = 0.005 | $R^2$ = 0.000 |
| Hippocampus | FAAH *vs* CB1R-47 | *APOE* -ε4 | *APOE* +ε4 | *APOE* -ε4 by sex | | *APOE* +ε4 by sex | |
| | | | | Male | Female | Male | Female |
| | | P = 0.893 | P = 0.524 | P = 0.494 | P = 0.843 | P = 0.3787 | P = 0.929 |
| | | $R^2$ = 0.001 | $R^2$ = 0.015 | $R^2$ = 0.098 | $R^2$ = 0.003 | $R^2$ = 0.098 | $R^2$ = 0.001 |
| Hippocampus | FAAH *vs* CB1R-37 | *APOE* -ε4 | *APOE* +ε4 | *APOE* -ε4 by sex | | *APOE* +ε4 by sex | |
| | | | | Male | Female | Male | Female |
| | | P = 0.117 | P = 0.728 | P = 0.327 | P = 0.228 | P = 0.950 | P = 0.361 |
| | | $R^2$ = 0.124 | $R^2$ = 0.005 | $R^2$ = 0.191 | $R^2$ = 0.119 | $R^2$ = 0.001 | $R^2$ = 0.060 |

**SUPPL Table 19: Linear regression (by *APOE* ε4 status): FAAH vs CB1R species within hippocampus. Significant (P ≤ 0.05) observations, if any, are identified in bold text.**

| | | | |
|---|---|---|---|
| Hippocampus | FAAH *vs* CB1R-60 | Male | Female |
| | | P = 0.625 | P = 0.110 |
| | | $R^2$ = 0.016 | $R^2$ = 0.092 |
| Hippocampus | FAAH *vs* CB1R-47 | Male | Female |
| | | P = 0.325 | P = 0.920 |
| | | $R^2$ = 0.065 | $R^2$ = 0.000 |
| Hippocampus | FAAH *vs* CB1R-37 | Male | Female |
| | | P = 0.422 | P = 0.076 |
| | | $R^2$ = 0.044 | $R^2$ = 0.108 |

**SUPPL Table 20: Linear regression (by sex alone): Association between CB1R species within hippocampus. Significant ($P \leq 0.05$) observations, if any, are identified in bold text.**

| Hippocampus | MAGL *vs* CB1R-60 | *APOE* -ε4 | *APOE* +ε4 | *APOE* -ε4 by sex | | *APOE* +ε4 by sex | |
|---|---|---|---|---|---|---|---|
| | | | | Male | Female | Male | Female |
| | | P = 0.563<br>$R^2$ = 0.018 | P = 0.440<br>$R^2$ = 0.023 | P = 0.396<br>$R^2$ = 0.147 | P = 0.318<br>$R^2$ = 0.083 | P = 0.760<br>$R^2$ = 0.010 | P = 0.290<br>$R^2$ = 0.080 |
| Hippocampus | MAGL *vs* CB1R-47 | *APOE* -ε4 | *APOE* +ε4 | *APOE* -ε4 by sex | | *APOE* +ε4 by sex | |
| | | | | Male | Female | Male | Female |
| | | P = 0.963<br>$R^2$ = 0.000 | P = 0.456<br>$R^2$ = 0.022 | P = 0.750<br>$R^2$ = 0.022 | P = 0.724<br>$R^2$ = 0.011 | **P = 0.002**<br>$R^2$ = 0.634 | P = 0.662<br>$R^2$ = 0.014 |
| Hippocampus | MAGL *vs* CB1R-37 | *APOE* -ε4 | *APOE* +ε4 | *APOE* -ε4 by sex | | *APOE* +ε4 by sex | |
| | | | | Male | Female | Male | Female |
| | | P = 0.934<br>$R^2$ = 0.000 | P = 0.225<br>$R^2$ = 0.056 | P = 0.451<br>$R^2$ = 0.118 | P = 0.876<br>$R^2$ = 0.002 | P = 0.544<br>$R^2$ = 0.038 | P = 0.301<br>$R^2$ = 0.076 |

**SUPPL Table 21: Linear regression (by *APOE* ε4 status): MAGL vs CB1R species within hippocampus. Significant (P ≤ 0.05) observations, if any, are identified in bold text.**

| Hippocampus | MAGL *vs* CB1R-60 | Male | Female |
|---|---|---|---|
| | | P = 0.550<br>$R^2$ = 0.021 | P = 0.748<br>$R^2$ = 0.004 |
| **Hippocampus** | **MAGL *vs* CB1R-47** | **Male** | **Female** |
| | | **P = 0.001**<br>$R^2$ = 0.489 | P = 0.541<br>$R^2$ = 0.014 |
| **Hippocampus** | **MAGL *vs* CB1R-37** | **Male** | **Female** |
| | | P = 0.554<br>$R^2$ = 0.021 | P = 0.488<br>$R^2$ = 0.018 |

**SUPPL Table 22: Association between MAGL and CB1R species within hippocampus (by sex alone). Significant (P ≤ 0.05) observations, if any, are identified in bold text.**

| Cortex *vs* Hippocampus | 60 *vs* 60 | Pooled | By males | By WT |
|---|---|---|---|---|
| | | P = 0.994 | P = 0.272 | P = 0.153 |
| | | $R^2 = 0.000$ | $R^2 = 0.196$ | $R^2 = 0.308$ |
| | | | By females | By Tg |
| | | | P = 0.827 | P = 0.437 |
| | | | $R^2 = 0.005$ | $R^2 = 0.104$ |
| Cortex *vs* Hippocampus | 47 *vs* 47 | Pooled | By males | By WT |
| | | P = 0.425 | P = 0.207 | P = 0.374 |
| | | $R^2 = 0.046$ | $R^2 = 0.250$ | $R^2 = 0.133$ |
| | | | By females | By Tg |
| | | | P = 0.232 | P = 0.767 |
| | | | $R^2 = 0.228$ | $R^2 = 0.016$ |
| Cortex *vs* Hippocampus | 37 *vs* 37 | Pooled | By males | By WT |
| | | **P = 0.006** | **P = 0.012** | P = 0.652 |
| | | $R^2 = 0.434$ | $R^2 = 0.681$ | $R^2 = 0.036$ |
| | | | By females | By Tg |
| | | | P = 0.166 | P = 0.454 |
| | | | $R^2 = 0.293$ | $R^2 = 0.097$ |

**SUPPL Table 23: Linear regression (by genotype): CB1R species between mouse cortex and hippocampus. Significant ($P \leq 0.05$) observations, if any, are identified in bold text.**

| Cortex | 60 *vs* 47 | Pooled | By males | By WT |
|---|---|---|---|---|
| | | P = 0.180 | P = 0.505 | P = 0.659 |
| | | $R^2$ = 0.125 | $R^2$ = 0.078 | $R^2$ = 0.035 |
| | | | By females | By Tg |
| | | | P = 0.178 | P = 0.112 |
| | | | $R^2$ = 0.278 | $R^2$ = 0.366 |
| Cortex | 60 *vs* 37 | Pooled | By males | By WT |
| | | P = 0.406 | P = 0.595 | P = 0.265 |
| | | $R^2$ = 0.050 | $R^2$ = 0.050 | $R^2$ = 0.202 |
| | | | By females | By Tg |
| | | | P = 0.647 | P = 0.742 |
| | | | $R^2$ = 0.037 | $R^2$ = 0.019 |
| Cortex | 47 *vs* 37 | Pooled | By males | By WT |
| | | P = 0.345 | **P = 0.035** | P = 0.827 |
| | | $R^2$ = 0.064 | $R^2$ = 0.552 | $R^2$ = 0.009 |
| | | | By females | By Tg |
| | | | P = 0.979 | P = 0.243 |
| | | | $R^2$ = 0.000 | $R^2$ = 0.218 |

**SUPPL Table 24: Linear regression (by genotype): CB1R species *vs* CB1R species within mouse cortex. Significant (P ≤ 0.05) observations, if any, are identified in bold text.**

| Hippocampus | 60 vs 47 | Pooled | By males | By WT |
|---|---|---|---|---|
| | | P = 0.297 | P = 0.255 | P = 0.375 |
| | | $R^2$ = 0.077 | $R^2$ = 0.209 | $R^2$ = 0.133 |
| | | | By females | By Tg |
| | | | P = 0.532 | **P = 0.031** |
| | | | $R^2$ = 0.068 | $R^2$ = 0.565 |
| Hippocampus | 60 vs 37 | Pooled | By males | By WT |
| | | P = 0.502 | P = 0.321 | P = 0.452 |
| | | $R^2$ = 0.033 | $R^2$ = 0.163 | $R^2$ = 0.097 |
| | | | By females | By Tg |
| | | | P = 0.737 | P = 0.204 |
| | | | $R^2$ = 0.020 | $R^2$ = 0.019 |
| Hippocampus | 47 vs 37 | Pooled | By males | By WT |
| | | **P < 0.0001** | **P < 0.0001** | **P = 0.003** |
| | | $R^2$ = 0.887 | $R^2$ = 0.961 | $R^2$ = 0.793 |
| | | | By females | By Tg |
| | | | **P = 0.040** | **P = 0.002** |
| | | | $R^2$ = 0.532 | $R^2$ = 0.824 |

**SUPPL Table 25: Linear regression (by genotype): CB1R species *vs* CB1R species within mouse hippocampus. Significant (P ≤ 0.05) observations, if any, are identified in bold text.**

| Hippocampus | FAAH *vs* CB1R-60 | Pooled | By males | By WT |
|---|---|---|---|---|
| | | P = 0.480 | P = 0.936 | P = 0.480 |
| | | $R^2$ = 0.036 | $R^2$ = 0.001 | $R^2$ = 0.086 |
| | | | By females | By Tg |
| | | | P = 0.553 | P = 0.652 |
| | | | $R^2$ = 0.036 | $R^2$ = 0.038 |
| Hippocampus | FAAH *vs* CB1R-47 | Pooled | By males | By WT |
| | | P = 0.110 | P = 0.317 | P = 0.203 |
| | | $R^2$ = 0.173 | $R^2$ = 0.165 | $R^2$ = 0.254 |
| | | | By females | By Tg |
| | | | P = 0.266 | P = 0.468 |
| | | | $R^2$ = 0.201 | $R^2$ = 0.091 |
| Hippocampus | FAAH *vs* CB1R-37 | Pooled | By males | By WT |
| | | P = 0.175 | P = 0.388 | P = 0.243 |
| | | $R^2$ = 0.128 | $R^2$ = 0.126 | $R^2$ = 0.218 |
| | | | By females | By Tg |
| | | | P = 0.126 | P = 0.679 |
| | | | $R^2$ = 0.000 | $R^2$ = 0.030 |

**SUUPL Table 26: Linear regression (by genotype): CB1R species *vs* FAAH within mouse hippocampus. Significant (P ≤ 0.05) observations, if any, are identified in bold text.**

| Hippocampus | MAGL *vs* CB1R-60 | Pooled | By males | By WT |
|---|---|---|---|---|
| | | P = 0.858 | P = 0.085 | **P = 0.044** |
| | | $R^2$ = 0.002 | $R^2$ = 0.415 | $R^2$ = 0.518 |
| | | | By females | By Tg |
| | | | P = 0.037 | P = 0.510 |
| | | | $R^2$ = 0.543 | $R^2$ = 0.076 |
| Hippocampus | MAGL *vs* CB1R-47 | Pooled | By males | By WT |
| | | P = 0.130 | P = 0.746 | P = 0.091 |
| | | $R^2$ = 0.156 | $R^2$ = 0.018 | $R^2$ = 0.403 |
| | | | By females | By Tg |
| | | | P = 0.337 | P = 0.660 |
| | | | $R^2$ = 0.154 | $R^2$ = 0.034 |
| Hippocampus | MAGL *vs* CB1R-37 | Pooled | By males | By WT |
| | | P = 0.066 | P = 0.586 | P = 0.157 |
| | | $R^2$ = 0.221 | $R^2$ = 0.052 | $R^2$ = 0.303 |
| | | | By females | By Tg |
| | | | P = 0.462 | P = 0.367 |
| | | | $R^2$ = 0.093 | $R^2$ = 0.137 |

**SUPPL Table 27: Linear regression (by genotype): CB1R species *vs* MAGL within mouse hippocampus. Significant (P ≤ 0.05) observations, if any, are identified in bold text.**
