## Supplementary material for "Differences in the co-distribution of the Cannabinoid Receptor-1, FAAH, and MAGL in the human and mouse brain could be limiting clinical translation of endocannabinoid outcomes in mouse models of Alzheimer disease": SUPPL Figures 1-4

SUPPL Figure 1: Mean expression levels of CB1Rs in either Cortex or Hippocampus stratified by *APOE* ε4 status alone

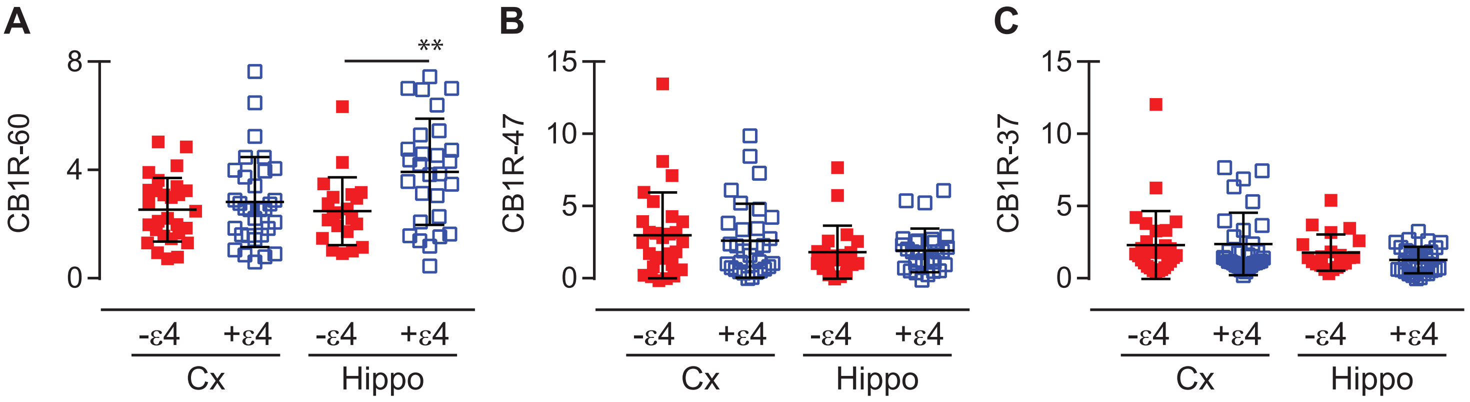

SUPPL Figure 2: Mean expression levels of CB1Rs in either Cortex or Hippocampus stratified by sex alone

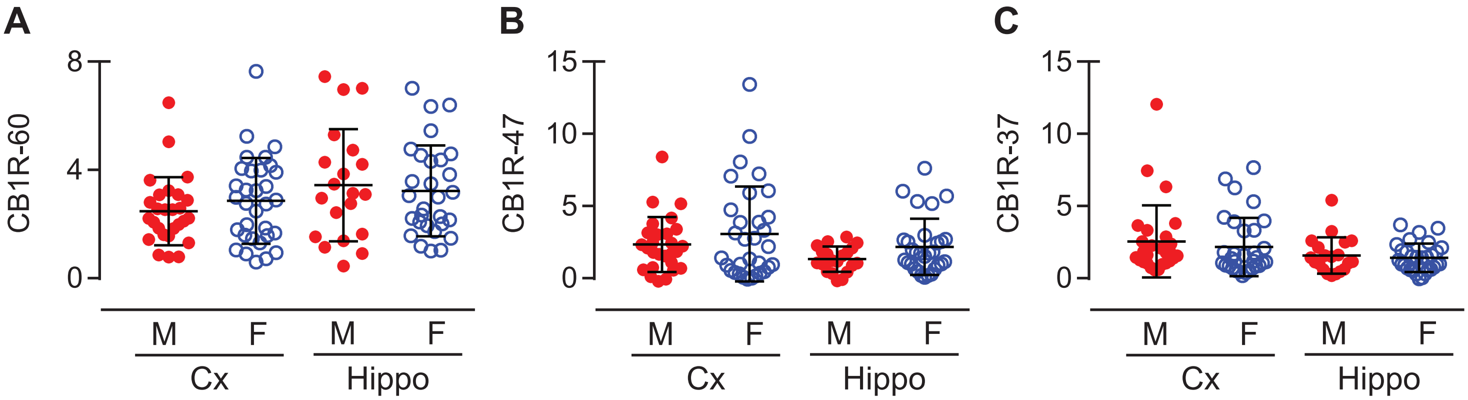

SUPPL Figure 3: Mean expression levels of CB1Rs in Hippocampus stratified by *APOE* ε4-by-sex (independent of diagnosis)

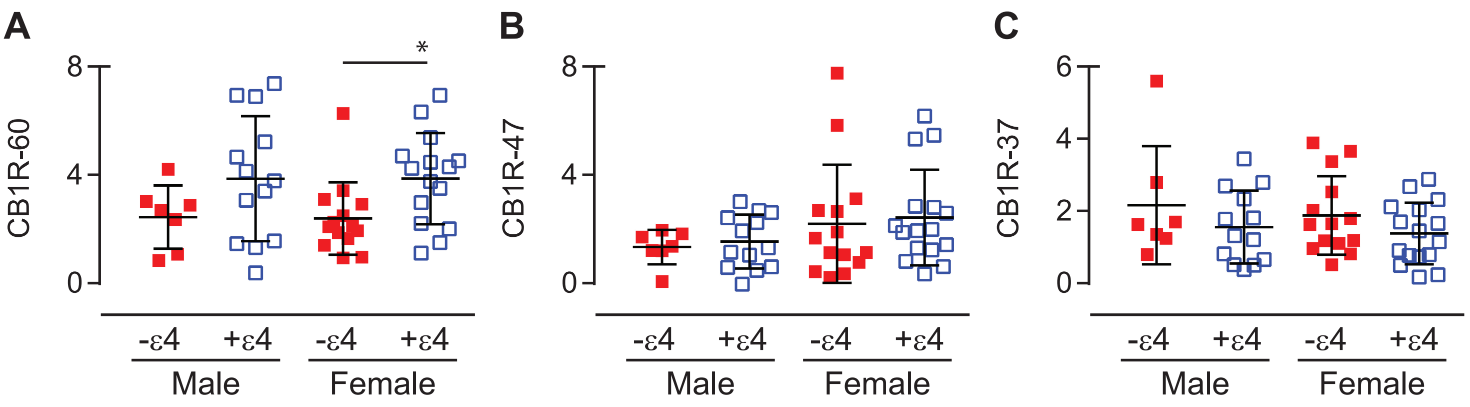

SUPPL Figure 4: Mean expression levels of FAAH and MAGL in either Cortex or Hippocampus stratified by *APOE* ε4 status alone or by sex alone

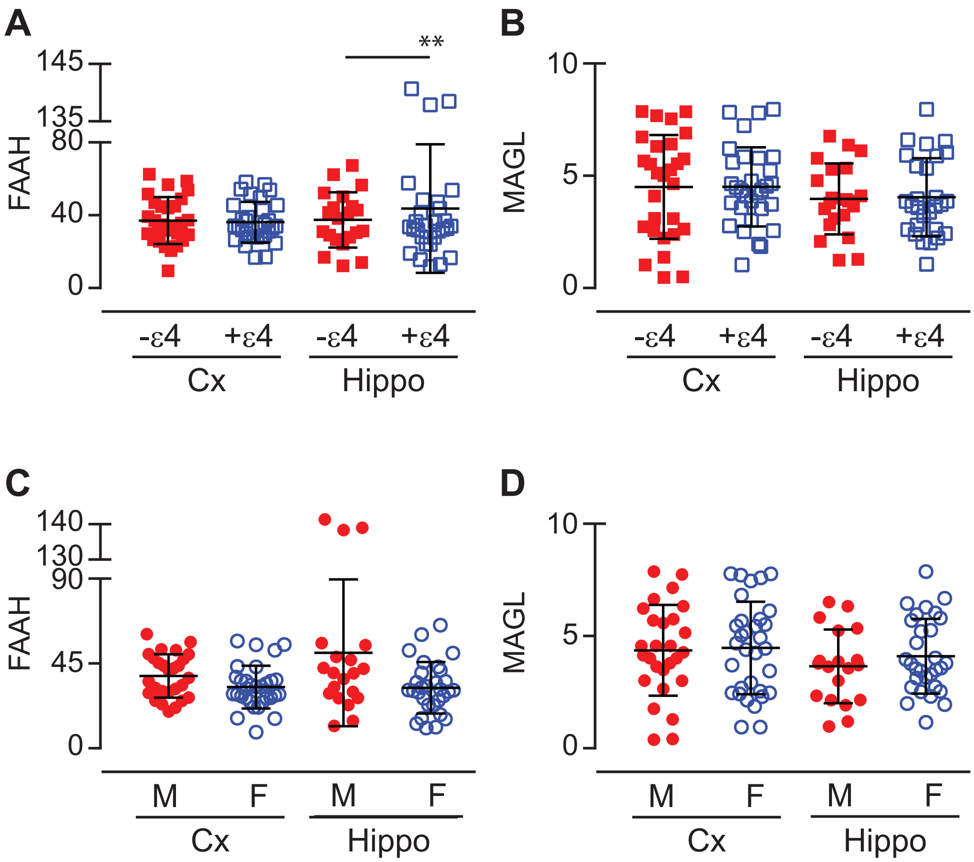
